# Real-time imaging of transcriptional feedback in nonsense-mediated mRNA decay

**DOI:** 10.1101/2025.05.20.655238

**Authors:** Md. Dobirul Islam, Tamoghna Das, Robert H. Singer, Hanae Sato

## Abstract

Nonsense-mediated mRNA decay (NMD) is a translation-coupled mRNA decay pathway triggered by a premature termination codon (PTC). While in-frame stop codons are typically defined by cytoplasmic ribosomes, unexpected changes in transcription have been reported in genes containing PTCs. This observation suggests the possibility of PTC detection at the transcription site, which has not been thoroughly investigated with high temporal and spatial resolution. Here, we utilize a real-time imaging approach to simultaneously detect transcription sites expressing wild-type or NMD-targeted β-globin reporter genes in the same cell. Our data indicates a dynamic change in the transcription of PTC-containing β-globin mRNA that depends on translation, NMD, and nuclear protein import, supporting the existence of rapid transcriptional feedback following NMD in the cytoplasm. This study establishes a robust temporal link between cytoplasmic mRNA decay and nuclear transcription.

**One-Sentence Summary:** Live-cell imaging shows that nonsense-mediated mRNA decay triggers a transcriptional feedback loop.

## Main Text

The flow of gene expression from transcription to translation, commonly known as the central dogma, is typically portrayed as a unidirectional process. However, several studies have provided evidence of crosstalk between cytoplasmic mRNA decay and transcription (*1*). Such crosstalk can occur through mechanisms that alter the rates of mRNA decay and transcription, thereby regulating cellular protein concentrations (*2–4*). Notably, decay-transcription feedback coupling has also been observed in mammalian cells, including during gamma-herpesvirus infection, where SOX-mediated cytoplasmic mRNA decay triggers transcriptional repression (*5*), and in nonsense-mediated mRNA decay (NMD) pathways (*6–9*).

NMD is a critical, evolutionarily conserved mRNA surveillance pathway in eukaryotes that prevents the production of harmful truncated proteins by eliminating mRNAs containing premature termination codons (PTCs)(*10*, *11*). It is initiated during translation termination, when release factors (eRF1/eRF3) recognize a PTC and recruit NMD core factors. UPF1 forms a surveillance complex at the PTC and, together with SMG1, UPF2, and UPF3, converts the aberrant termination event into targeted mRNA degradation (*12–14*).

In mammals, a downstream exon-junction complex (EJC) helps mark a stop codon as premature (*15*, *16*), while unusually long 3′ UTRs can also trigger NMD by weakening interactions between PABPC1 and termination factors (the Faux 3’UTR model) (*17–19*), making 3’UTR length an important determinant of decay. Once activated, NMD degrades transcripts through SMG6-mediated cleavage (*20*, *21*) or SMG7-mediated exonucleolytic decay (*22*), and the redundancy of these pathways highlights the robustness of this surveillance system (*23*, *24*).

In contrast to the global regulation of transcription-decay rates, the transcriptional changes of PTC-containing transcripts are sequence-specific, with these transcripts modulating the expression of their own or homologous genes. Specifically, PTC-containing pre-mRNAs, but not those with missense or frameshift mutations, have been found to accumulate at the transcription sites of immunoglobulin (Ig) µ and T cell receptor (TCR) β genes (*6*, *9*). This accumulation results from transcriptional retention rather than increased transcriptional activity.

This PTC-specific feedback highlights transcriptional adaptation, an emerging phenomenon in which the deleterious effects of nonsense mutations are counteracted by the upregulation of homologous genes, although the underlying mechanism remains largely unknown (*7*, *8*, *25*, *26*). This phenomenon is now recognized as a conserved and widespread biological mechanism that buffers genetic perturbations and maintains physiological function. Notably, recent studies have highlighted its therapeutic potential to restore gene function through the compensatory activation of functionally related or paralogous genes (*26*). These insights underscore the critical role of NMD-triggered transcriptional regulation in genetic compensation, revealing nonsense-associated transcriptional regulation as both a fundamental principle of gene expression control and a promising avenue for therapeutic intervention.

Moreover, in cells carrying heterozygous disease-associated nonsense mutations, the levels of normal mRNA expressed from the wild-type allele were found to be reduced (*27*, *28*), suggesting that nonsense-associated transcriptional alterations may influence the severity of phenotypic expression in heterozygous nonsense mutations. These findings raise important concerns about the broader regulatory consequences of nonsense mutations beyond the mutant transcript itself. Despite these emerging insights, the molecular mechanisms by which cytoplasmic mRNA decay influences nuclear transcription remain largely elusive, representing a critical gap in our understanding of gene expression regulation and its disruption in genetic disease.

To investigate the relationship between transcriptional activity and cytoplasmic NMD, we developed a real-time imaging system to monitor the transcriptional dynamics of an NMD reporter gene in living cells. This system utilizes a ponasterone A (ponA)-inducible bidirectional reporter that co-expresses both PTC-free and PTC-containing β-globin mRNAs, each tagged with MS2 or PP7 in their 3’UTRs (*29*) (Fig. 1A). By enabling simultaneous detection of transcription in living cells, our system allows for precise examination of the dynamic transcriptional response during mRNA decay with high spatiotemporal resolution.

**Fig. 1.**
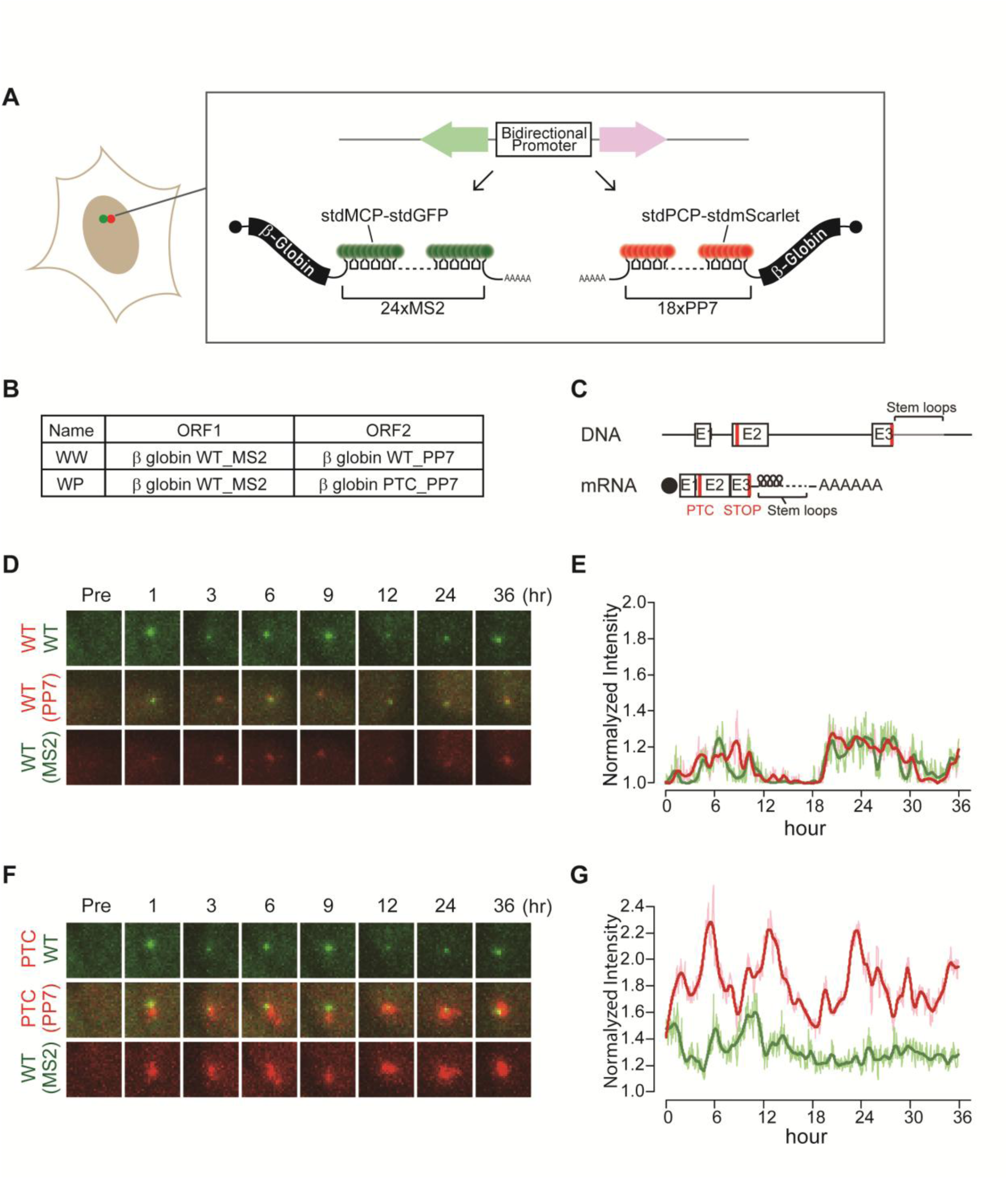
Simultaneous real-time tracking of transcription activities at the NMD reporter with and without PTC. **(A)** Schematic of PonA-inducible bi-directional promoter expressing NMD reporter β-globin genes. The transcripts expressed from either direction of the promoter contain MS2 or PP7 sequences in the 3’UTR, which are labeled with stdMCP-stdGFP or stdPCP-stdmScarlet. 24 or 18 repeats of MS2 or PP7, resulting in a similar 3’UTR length, were used. **(B)** The table shows the mRNAs expressed from the bi-directional promoter of each construct (WW; Wild-type and Wild-type or WP; Wild-type and PTC-containing β-globin). WT; wild-type, PTC; premature termination codon. **(C)** Structures of the NMD reporter Gl (DNA; upper and mRNA; lower) containing a PTC at position 39 (PTC). Horizontal lines represent introns, 5’UTR and 3’UTR, while boxes represent each of the three Gl exons (E1-3) joined by splicing-generated exon-exon junctions. Black dot represents the cap structure; red lines indicate termination codons; STOP represents normal termination codon; AAAAAA indicates poly(A) tail. **(D-G)** Images of transcription sites before (Pre) and after (1-36 hours) transcription induction by PonA. The image size of each transcription site shown here is 7 x 7 µm². **(E & G)** Normalized intensity of transcription sites was plotted. Each transcription site was imaged every 2 minutes, and the intensities of transcription sites were detected using TrackMate (*1*). Imaging settings were kept the same for all constructs. The mean intensity of the transcription site was normalized by the mean nuclear intensity at each time point. The mean intensity of the transcription site was normalized by dividing the spot intensity by the mean nuclear intensity at each time point. Normalized intensity at each transcription site of Gl-MS2 (green) or Gl-PP7 (pink) was smoothed and fitted using the lowess method (Gl-MS2 (dark green) or Gl-PP7 (red)) in R.

To establish a control, we expressed the wild-type β-globin gene in both directions within a single cell (Fig. 1B; WW: Wild-type and Wild-type). For the NMD reporter, we expressed wild-type and PTC-containing β-globin genes in opposite orientations within the same cell (Fig. 1B and C, WP: Wild-type and PTC). U2OS cells stably expressing WW or WP were generated using the Flp-In system, enabling us to compare transcriptional activity at the same chromatin locus while minimizing variability associated with transcription at different chromatin locations (*29*, *30*). Given that the insertion of multiple stem loops in the 3’UTR could potentially induce NMD (*31*), we assessed the impact of a 3’UTR extension on mRNA stability by inserting stem loops and knocking down UPF1 using shRNA. RT-qPCR analysis of endogenous UPF1 and the control β-actin mRNA confirmed efficient UPF1 knockdown (Fig. S1A), which led to an increased abundance of endogenous NMD target transcripts, indicating effective inhibition of NMD (Fig. S1B). Moreover, comparison of reporter mRNA levels containing MS2 or PP7 stem loops in the WW construct between control shRNA– and UPF1 shRNA–expressing cells showed that insertion of these stem loops did not elicit NMD in this system (Fig. S1C and D). We further confirmed, using the WP construct, that UPF1 knockdown resulted in a marked increase in the PTC-containing PP7 transcript, demonstrating that the PTC-containing PP7 transcript is subject to NMD (Fig. S2A and B).

To simultaneously examine transcriptional dynamics at wild-type and PTC-containing mRNA transcription sites, we imaged transcriptional activity in cells expressing either the WW (Fig. 1D and E) or WP (Fig. 1F and G) every two minutes in real-time. Following transcription induction with PonA, fluorescence intensities at transcription sites were quantified using TrackMate (*32*). The analytical pipeline used to identify transcriptional bursts (peaks) in Figure 1 is outlined in Figure S3A. We then analyzed the average intensities at peaks in multiple cells expressing WW (Fig. S3B) or WP (Fig. S3C). Notably, we observed enhanced transcriptional bursts in WP-expressing cells (Fig. 1F and G, Fig. S3C, movie S2) compared to WW-expressing cells (Fig. 1D and E, Fig. S3B, movie S1) over a 36-hour period. Because we observed time-dependent changes in size of transcription site (TS-area), we next assessed whether enlargement of TS-area influenced TrackMate-based detection. Although TS-area measurements rely on threshold-based masking and therefore do not define precise boundaries, TS area showed a positive correlation with TrackMate-detected TS fluorescence signals (Fig.S3D). These results indicate that TS enlargement is reflected in the TrackMate-based TS detection and does not account for the observed differences in transcriptional bursting between WW and WP constructs.

The transcription site in the WW construct, which expresses wild-type mRNA from both directions of the promoter, appeared as compact fluorescent spots (Fig. 2A). In contrast, the transcription site in the WP construct, which expresses wild-type and PTC-containing mRNAs, exhibited pronounced dynamic changes in shape and size and was frequently observed as a stretched or enlarged structure at the PTC-containing allele (Fig. 2B). This phenotype was consistently observed with both GFP-and mScarlet-tagged reporters (Fig. 2B).

**Fig. 2.**
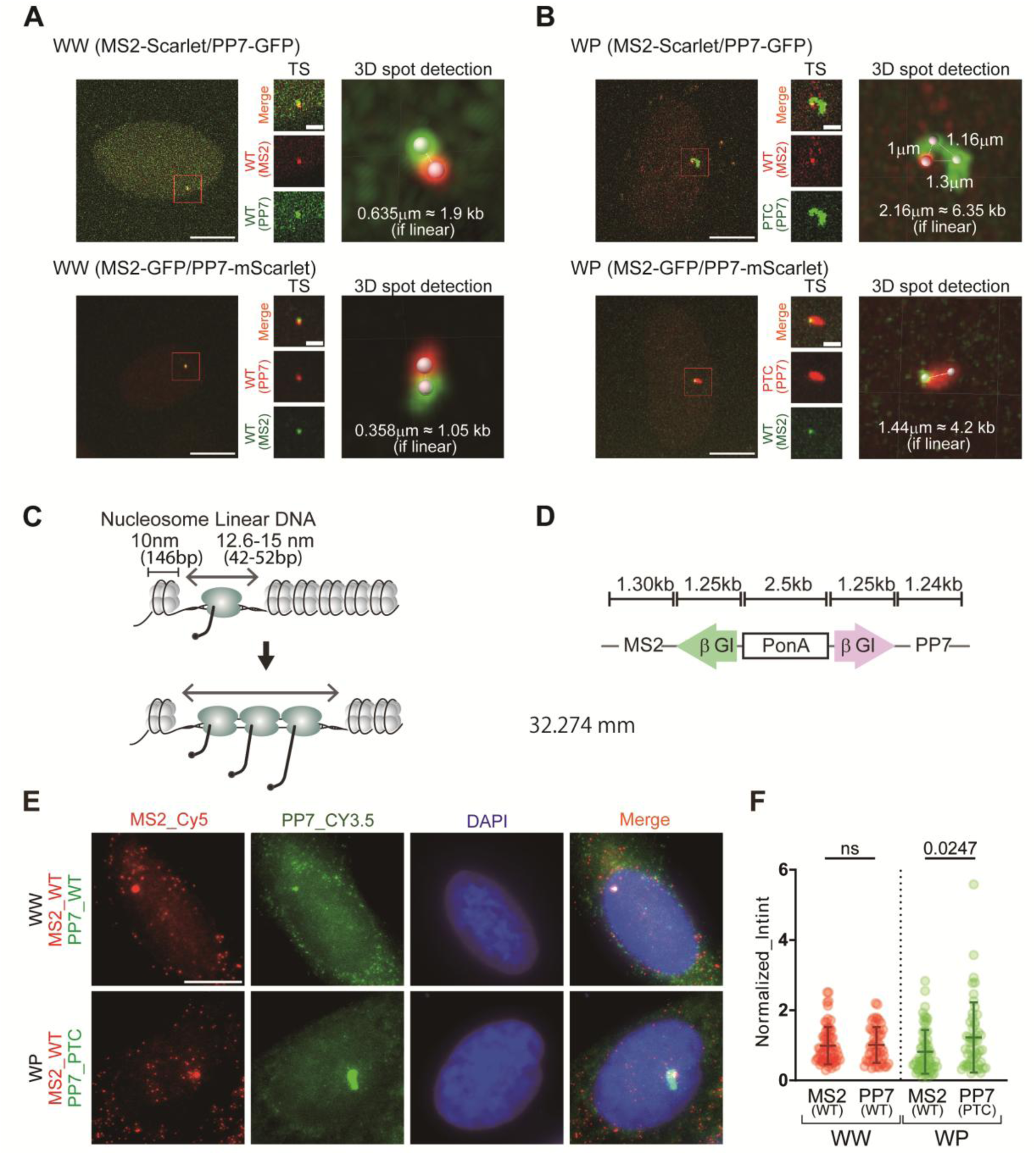
Simultaneous detection of transcription sites with and without a PTC reveals PTC allele-specific transcription-site enlargement. **(A)** Image of a transcription site expressing wild-type β-globin mRNA in both orientations (WW). **(B)** Images of transcription sites expressing either wild-type or PTC-containing β-globin mRNA from each orientation of the bidirectional promoter (WP). Scale bar = 10 µm (left). Insets show higher magnification. Scale bar = 2 µm (middle). The region outlined in red is shown on the right of each image. The physical distance between two transcription sites was measured from fluorescence images as the center-to-center distance of the spots using Imaris viewer. **(C)** Schematic representation of stretched chromatin structure due to occupancy by elongating RNA polymerase II. Nucleosome-free DNA is assumed to extend 12.6–15 nm per 42–52 bp, corresponding to approximately 0.34 µm per 1 kb, based on previous measurements. **(D)** Approximate length of the reporter construct. **(E)** Detection of transcription sites expressing WW or WP by smFISH. Scale bar, 10 µm. **(F)** Quantification of transcription sites in cells expressing WW or WP detected by smFISH. Nascent transcript detection was performed using FISH-quant to determine the integrated intensity (IntInt) at each transcription site. Integrated intensities were normalized to the mean integrated intensity across the entire cell population of WW and WP for each fluorophore (Normalized_IntInt). Data was obtained from 69 (WW) and 64 (WP) cells. Statistical analysis was performed using two-way ANOVA in GraphPad Prism (ns, not significant).

Using confocal microscopy combined with deconvolution and 3D spot detection, we estimated the apparent spatial separation between the two bidirectionally transcribed TSs (TS-distance). In the WW construct, this distance ranged from ∼0.35 to 0.7 μm. In contrast, the WP construct exhibited TS-distance of up to ∼2.2 μm (approximately 6.4 kb assuming linear DNA), with the extent of separation varying with transcriptional bursting activity (Fig. 2B–D), suggesting that transcription of the PTC-containing locus involves high RNA polymerase II occupancy and chromatin stretching (*33*). Notably, the genomic length of the GI-PP7 gene is approximately 2.5 kb, which is shorter than the estimated spatial extension (∼4kb). This discrepancy may arise from differences in local chromatin organizations associated with the PonA promoter, diffusion-related overestimation of fluorescent spot centroids, or other currently unknown mechanisms. Importantly, a similar PTC-specific TS enlargement was also observed for another NMD substrate, the PTC-containing Igμ mini-gene (Fig. S4), supporting the generality of this phenomenon in NMD. This PTC-specific TS enlargement was further confirmed by nascent transcript detection using single-molecule fluorescence in situ hybridization (smFISH) (Fig 2E and F). The integrated fluorescence intensity at the PTC-containing transcription site was increased; however, the extent of TS enlargement was variable, likely reflecting the stochastic nature of transcriptional bursting, which does not always capture the enlarged state at the time of fixation.

Enlarged transcription sites may arise from increased transcription initiation, frequent pausing, slower RNA polymerase II (Pol II) progression, or impaired transcription termination, all of which may lead to stalled RNA polymerases. To investigate this further, we performed a pulse-chase assay, where transcription was induced with PonA which was then subsequently removed (Fig. 3A, B and C). Our results showed that while the fluorescence intensity at the transcription site of wild-type mRNA disappeared after PonA removal, the fluorescence intensity at the transcription site of PTC-containing mRNA persisted for over 100 minutes after inducer removal in WP (Fig. 3B, C and S5). To further examine RNA polymerase II (Pol II) occupancy, we performed Pol II-ChIP qPCR. Because the bidirectionally expressed genes share largely overlapping sequences, we designed primers targeting a chromatin region containing a unique sequence (Fig. 3D), enabling allele-specific detection (Fig. 3E and F). This analysis revealed increased Pol II association with the PTC-containing allele (Fig. 3E and F).

**Fig. 3.**
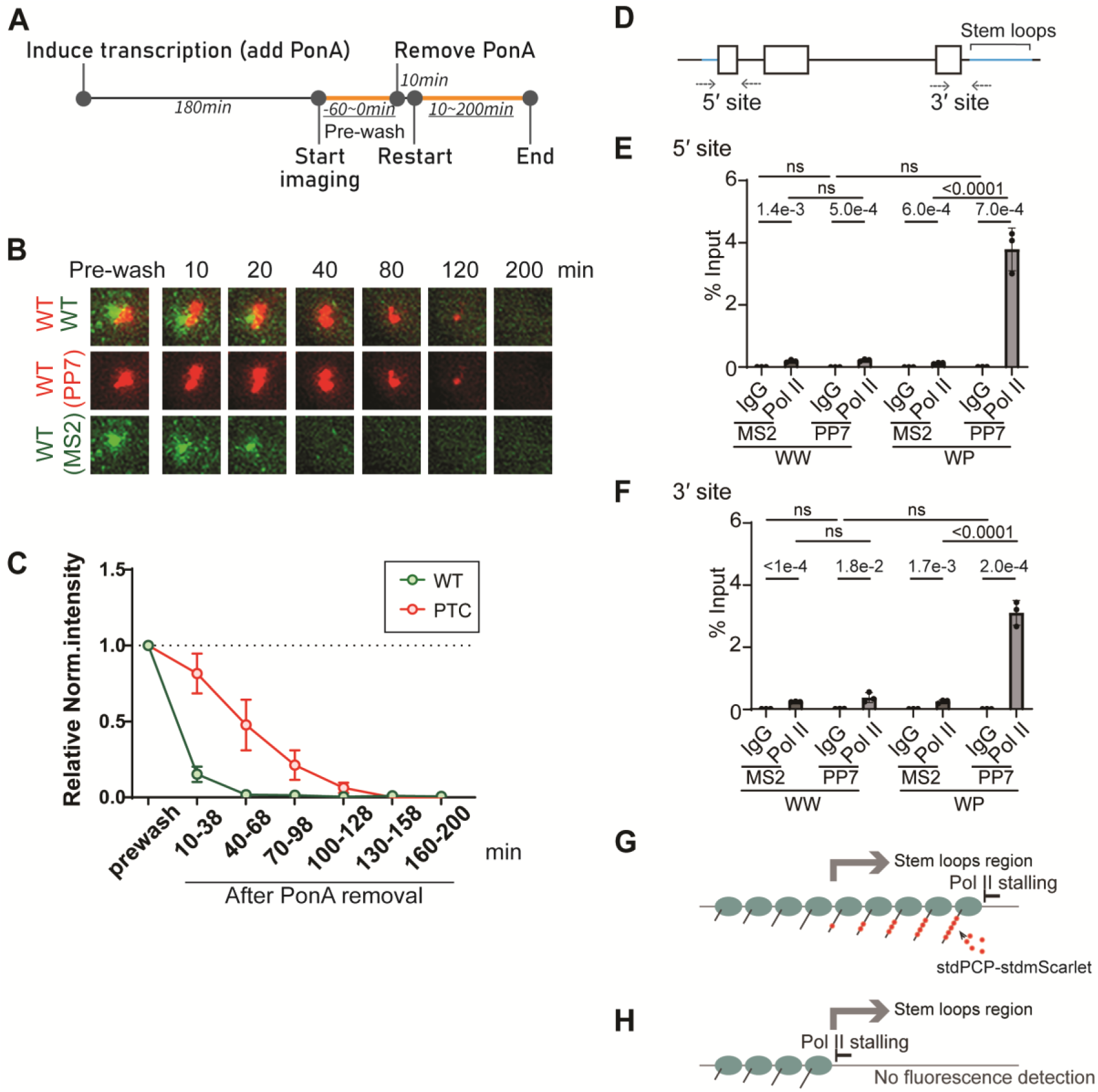
PTC allele–specific transcription-site enlargement is associated with RNA polymerase II stalling. **(A)** Detection of TS expressing the WP construct before and after removal of the transcription inducer. Timeline indicates transcriptional activation via PonA addition followed by its removal. Orange lines and underlined time points indicate the duration of real-time imaging. **(B)** Images of transcription sites before (Pre-wash) and after PonA removal (10–100 min). Each transcription site image represents a 7 × 7 µm² field. **(C)** Quantification of transcriptional activity from the WP construct before (Pre-wash) and after PonA removal (10–100 min). Dots represent the mean normalized intensity of transcription sites across 11 cells. Error bars represent the standard deviation within the cell population. Statistical analysis was performed using an unpaired, two-tailed *t*-test in GraphPad Prism. **(D)** Positions of primer pairs annealing to the reporter DNA. Because the two alleles are bidirectionally expressed from the same promoter and contain limited unique sequence (blue line), primer pairs were designed to anneal to allele-specific regions to distinguish MS2-and PP7-tagged transcripts. (E, F) RNA polymerase II (Pol II) enrichment at the 5′ site **(E)** and 3′ site **(F)**, detected by chromatin immunoprecipitation against RNA polymerase II followed by qPCR (Pol II ChIP–qPCR). ChIP–qPCR signals are shown as percent input. Statistical analysis was performed using two-way ANOVA with cell line (WW and WP) and genomic locus as factors (ns, not significant). Post hoc multiple comparisons were performed using Tukey’s test. Data represent mean ± SD from *n* biological replicates. IgG served as a negative control. (G, H) Schematics illustrating transcriptional stalling of elongating RNA polymerase II at the end **(G)** or in the middle **(H)** of the gene.

This elevated Pol II occupancy likely reflects stalled RNA polymerase II at the 3’end of the gene, downstream of the stem loops (Fig. 3G), as transcripts from ongoing transcription upstream of the stem loops would not be visible (Fig. 3H).

PTC-dependent transcriptional enlargement raises the question of how a PTC could influence the transcription site before mRNA is exported to the cytoplasm, given that current biological evidence suggests PTCs are recognized only by translating ribosomes in the cytoplasm. Possible mechanisms include recognition of the PTC during transcription or the presence of an unknown feedback signal from the cytoplasm to the transcription site following PTC recognition. To explore this, we examined transcription sites immediately after PonA induction, at which point mRNAs have not yet been exported or translated. Fluorescence intensities at transcription sites were measured every 2 minutes for the first 2.5 hours after induction in cells expressing either the WW (Fig. S6A) or WP (Fig. S6B) construct, and the average intensity of the transcription sites was calculated. The data showed that transcriptional enlargement occurred in WP-expressing cells, but not in WW-expressing cells, beginning approximately 1 to 1.5 hours after transcription initiation. This suggests that the enlargement is not triggered immediately by PTC recognition during transcription. Instead, the observed delay of 1 to 1.5 hours may reflect the time required for downstream processes such as mRNA export, translation, NMD, and subsequent feedback to the transcription site.

We next investigated whether the transcriptional effect is translation-dependent, as translation is required for PTC recognition by ribosomes and subsequent NMD activation in the cytoplasm. To test this, we used the translation inhibitor cycloheximide (CHX). First, we confirmed that CHX had no effect on transcription sites in cells expressing the WW construct (Fig. S7A). We then verified that NMD was efficiently inhibited by monitoring the stabilization of endogenous NMD target transcripts (Fig. S2C). Under these conditions, we examined whether translation inhibition affected the spatial expansion in PTC containing transcription sites. Our results showed that transcription site expansion was observed without CHX (Fig. S6B) but not in CHX-treated PTC containing transcription site (Fig. S7B). Furthermore, we found that increased transcript accumulation at the transcription site was abolished during CHX treatment but reappeared upon CHX removal (Fig. 4A-C), indicating that the feedback to the transcription site is translation-dependent.

**Fig. 4.**
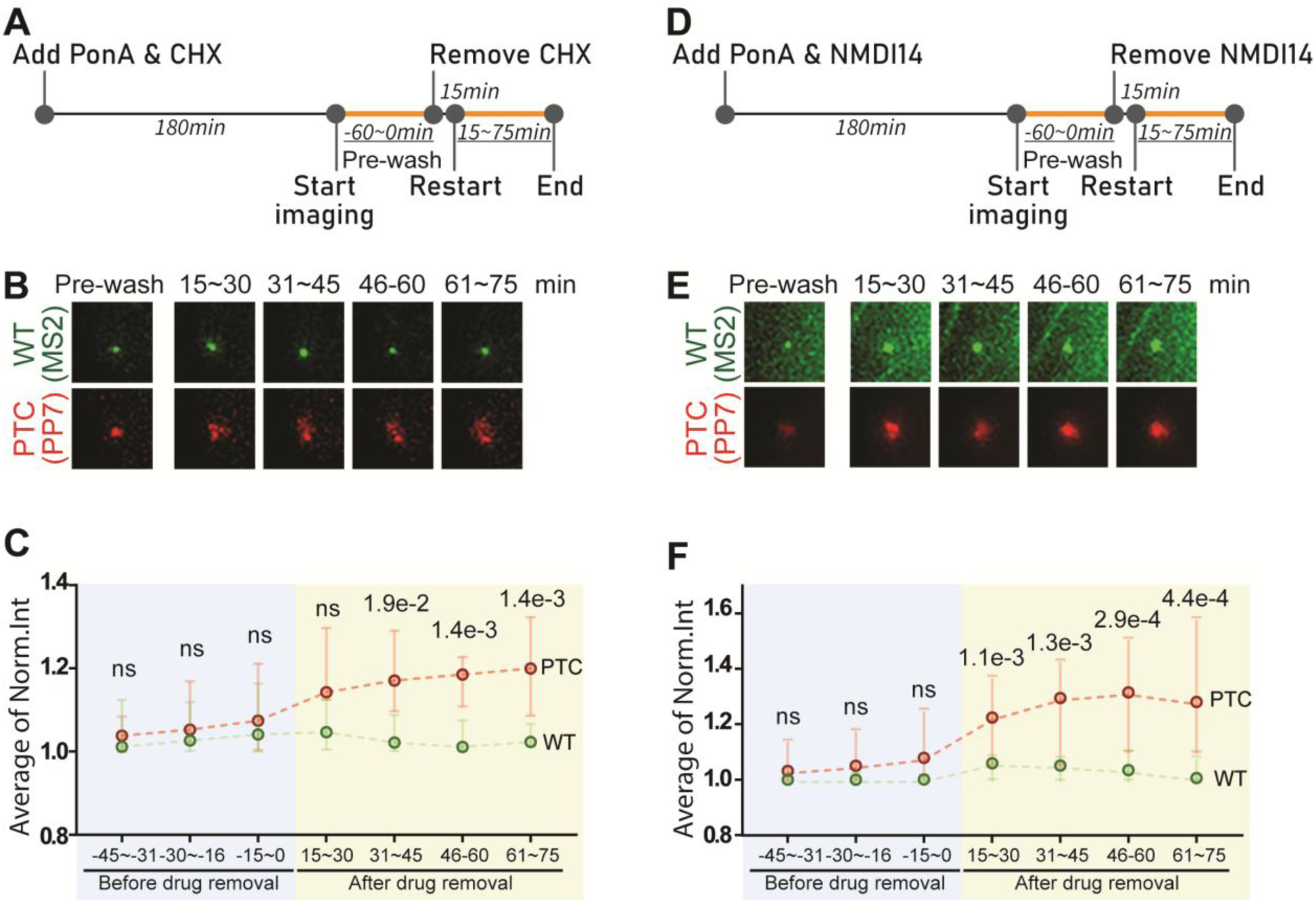
Rapid feedback of transcription preceded by translation and NMD. **(A)** Timeline of transcription induction by supplementation of PonA with CHX (100 µg/ml) followed by drug removal. Orange lines and underlined times indicate the time duration for real-time imaging. **(B)** Images of transcription site before (Pre-wash) and after removal of CHX (15-75 min). The image size of each transcription site shown here is 8 x 8 µm². **(C)** Detection of transcription sites expressing the WP construct before (Pre-wash) and after removal of CHX. Single points with 95% confidence intervals indicate the median normalized intensity for wild-type (green) and PTC-containing β-globin transcription sites (red) across 21 cells at each indicated time window. Statistical analysis was performed in GraphPad Prism using paired, nonparametric Wilcoxon matched-pairs signed-rank tests (ns, not significant). **(D)** Timeline of transcription induction by supplementation of PonA with the NMD inhibitor, NMDI14, followed by the removal of NMDI14 (40 µM). Orange lines and underlined times indicate the time duration for real-time imaging. **(E)** Images of transcription site before (Pre-wash) and after removal of NMDI14 (15-75 min). The image size of each transcription site shown here is 8 x 8 µm². **(F)** Detection of transcription sites expressing the WP construct before (Pre-wash) and after removal of NMDI14. Single points with 95% confidence intervals indicate the median normalized intensity for wild-type (green) and PTC-containing β-globin transcription sites (red) across 23 cells at each indicated time window. Statistical analysis was performed in GraphPad Prism using nonparametric Mann–Whitney tests without multiple-comparison correction (ns, not significant).

To further explore the role of NMD, we used the NMD inhibitor NMDI14, which interfere SMG7-UPF1 interactions (*34*). Similar to CHX, NMD was efficiently inhibited by NMDI14, as confirmed by stabilization of endogenous NMD target transcripts (Fig. S2C). Under these conditions, NMDI14 treatment blocked transcription-site enlargement, and its removal led to a rapid reappearance of the enlarged transcription site (Fig. 4D–F), further supporting that this phenomenon is NMD-dependent. Importantly, this effect was not due to medium replacement itself, as control experiments using DMSO-containing medium instead of NMDI14 had no impact on the transcription site (Figure S8).

The alteration of the transcription site was specific to the presence of the PTC, which resulted from a single nucleotide substitution in the wild-type gene. This sequence-specific effect suggests the existence of a mechanism that discriminates between mutated alleles in DNA or nascent mRNA at the transcription site. One possible explanation for the PTC-specific effect is the recognition of the allele by mRNA degradation fragments, which result from endonucleolytic cleavage of the mRNA near the PTC by SMG6 (*21*, *20*). These RNA fragments could potentially anneal with the DNA where the PTC-containing transcripts are originally produced. To test the role of mRNA fragments in transcriptional feedback, we introduced self-endonucleolytic mRNA cleavage by inserting a hammerhead ribozyme structure between the β-globin coding region and PP7 stem loop sequence in the WW construct (Fig. 5A; WH: Wild-type and Hammerhead). We generated a single-locus integrated WH construct in the PonA cell line and monitored transcriptional activity every two minutes for 36 hours. Unexpectedly, we observed comparable transcription activities (without enlargement) in WH-expressing cells, indicating that mRNA degradation fragments were not sufficient to trigger transcriptional feedback (Fig. 5B). However, we cannot exclude the possibility that the self-endonucleolytic mRNA cleavage by the hammerhead ribozyme failed to trigger transcriptional feedback due to insufficient cleavage efficiency, approximately 50% compared to the wild-type construct without the hammerhead ribozyme insertion, as confirmed by RT-qPCR (Fig. 5C). It is also possible that hammerhead ribozyme-mediated transcription feedback and its efficiency may be cell-type specific as previously reported in transcription adaptation (*26*).

**Fig. 5.**
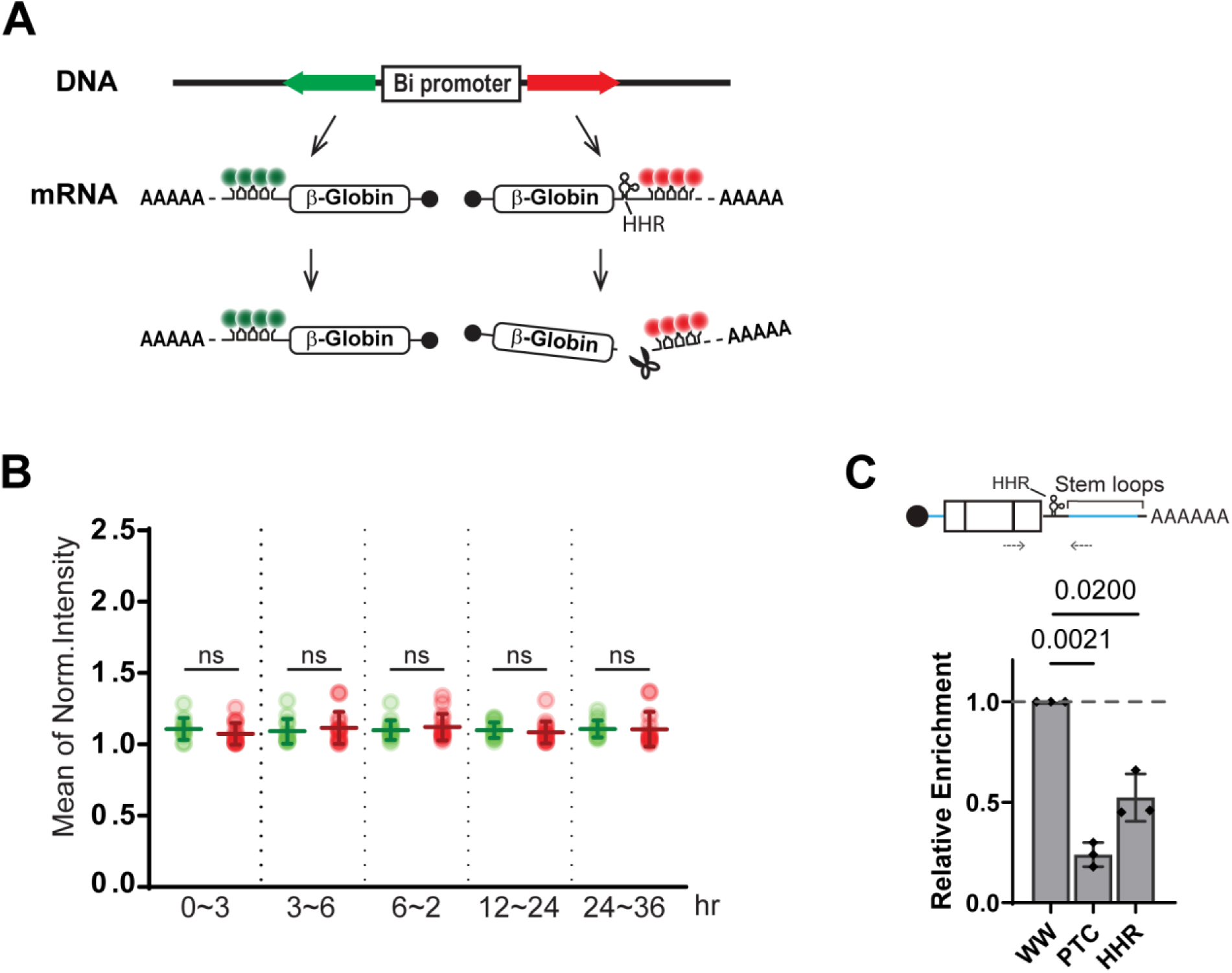
mRNA self-cleavage with hammerhead ribozyme (HHR) does not trigger transcriptional feedback. **(A)** Schematic of PonA inducible bi-directional promoter expressing wild-type β-globin genes with MS2 stem loops or with the self-cleaving motif of hammerhead ribozyme (HHR) between the β-globin ORF and PP7 stem loops in the 3’UTR that labeled with stdMCP-stdGFP or stdPCP-stdmScarlet. **(B)** Simultaneous detection of transcription sites expressing wild-type with (Red) or without HHR (Green). Single dots denote the mean of normalized intensity of transcription sites in single cells during the indicated time duration. P values were determined using two-tailed unpaired t-tests from 15 cells (ns, not significant). Error bars = Standard deviation in cell populations. **(C)** Quantitative detection of β globin mRNA with PP7 stem loops (Wild-type, PTC or HHR containing β globin expressing cells) were determined by RT-qPCR. MS2-tagged β globin mRNA (wild-type β globin mRNA in each cell) was used as a control. Error bars = Standard deviation from three independent experiments. The statistical analysis was performed by Graphpad prism software. The positions of primer pairs are illustrated above the graph.

Since self-endonucleolytic mRNA cleavage with the hammerhead ribozyme did not activate transcriptional feedback, it is likely that feedback is NMD-dependent, or RNA-binding proteins involved in mRNA decay are required for this mechanism. To investigate this possibility, we tested whether inhibition of nuclear protein import using the transport receptor importin-β inhibitor, importazole (*41*), reduces the transcriptional enlargement of the PTC-containing β-globin. We induced transcription under CHX treatment, then replaced CHX with fresh medium containing 20 µM PonA and 50 µM importazole (Fig. S9A and B). We monitored the transcription sites before and after the removal of CHX and the addition of importazole. The imaging data showed that the nuclear import inhibitor reduced the transcriptional enlargement of the PTC-containing transcript (Fig. S9A and B), suggesting the involvement of a protein or RNP regulator.

Notably, transcriptional feedback was observed exclusively at sites expressing PTC-containing transcripts, but not at sites expressing the wild-type transcripts, which were transcribed from the opposite side of a bi-directional promoter in this reporter system. In this setup, the key differences between the wild-type and PTC-containing transcripts lie in the presence of the PTC, the positioning of MS2 or PP7 stem loops within the 3’UTR, and the transcriptional orientation (positive vs. negative strand). We speculate that the limited sequence similarity in the 3’UTR, particularly at the sites of stem loop insertion, may underlie the observed allele-specific feedback. Specifically, the PTC-containing transcripts generate a PP7-tagged 3’fragment that is absent from the wild-type, potentially driving feedback at the transcription site (Fig. 6A). Based on this, we hypothesized that transcriptional feedback could occur at both wild-type and PTC-containing transcription sites if the 3’fragments generated during NMD share a common sequence context (Fig. 6B). To test this hypothesis, we inserted 6×MS2 or 5×PP7 stem loops into exon 1 of the β-globin gene, upstream of the PTC, while restoring the native 3’UTR (WWex and WPex constructs; Fig. 6A-D) and the resulting PTC-containing WPex transcript is subject to NMD (Fig. S2D). In contrast to constructs with stem loops in the 3’UTR, which generate 3’fragments with distinct downstream sequences in each allele (Fig. 6A), insertion of stem loops into exon 1 positions the cleavage site near the PTC and is therefore expected to generate 3’fragments with identical downstream sequences (Fig. 6B and C) (*21*, *20*). Remarkably, imaging analysis revealed transcriptional feedback at both wild-type (no PTC) and PTC-containing transcription sites under these conditions (Fig. 6D and E), supporting our hypothesis.

**Fig. 6.**
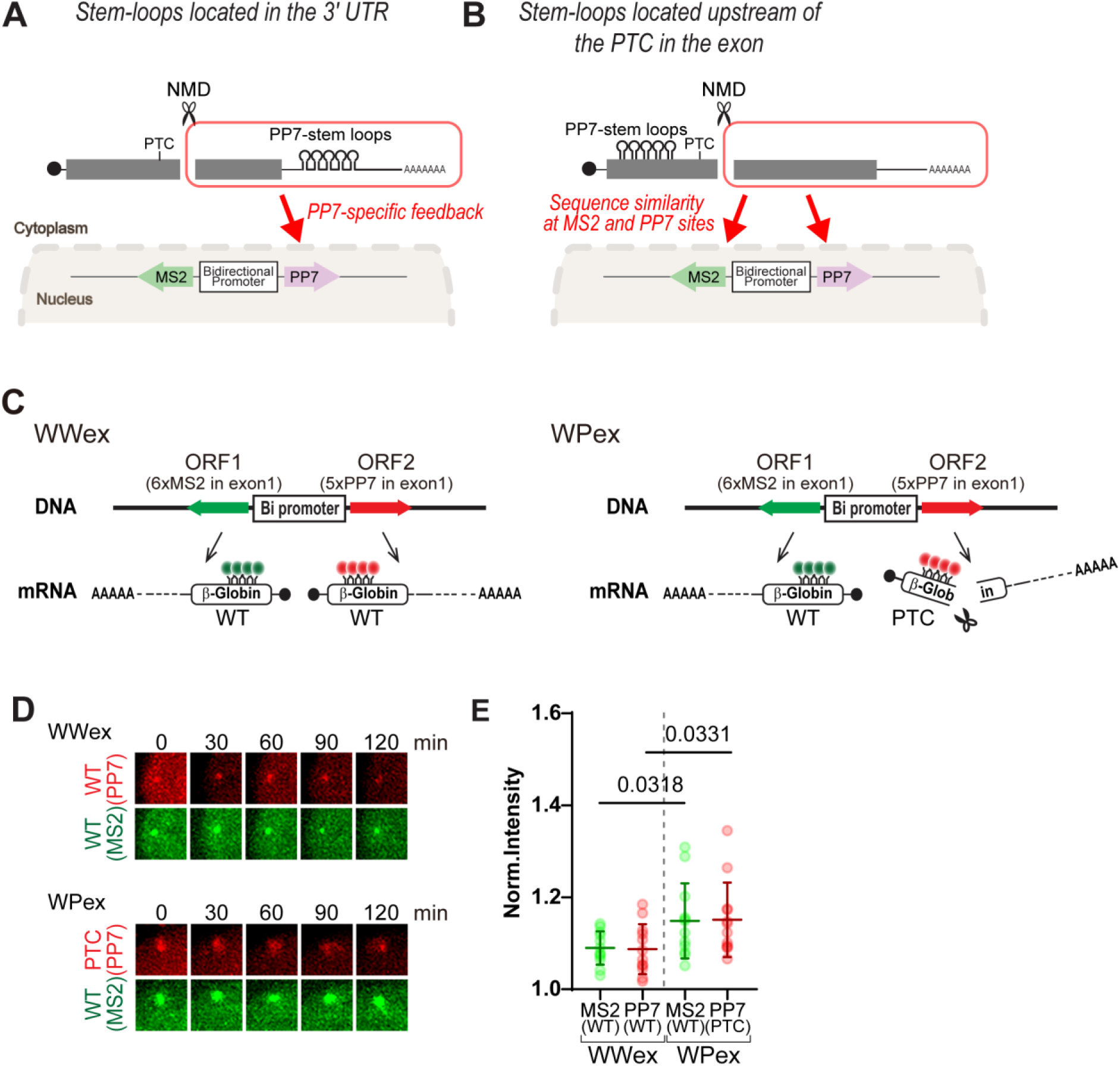
Insertion of stem loops upstream of the PTC within an exon enables transcriptional feedback in both alleles. **(A)** Schematic representation of stem-loop insertion into the 3’UTR. The 3’fragments generated by endonucleolytic cleavage during NMD provide feedback only to the PP7 allele due to sequence similarity in the 3’UTR. **(B)** Schematic representation of stem-loop insertion upstream of the PTC within an exon 1 and the expected feedback mechanism. The 3’ fragments generated by NMD cleavage can provide feedback to both the MS2 and PP7 alleles due to the shared sequence in the 3’UTR. **(C)** Schematic of the PonA-inducible bidirectional promoter driving expression of NMD reporter β-globin genes containing MS2 or PP7 stem-loop sequences in exon 1 in WWex (WT + WT with exon 1 stem loops) and WPex (WT + PTC with exon 1 stem loops), detected using stdMCP–stdGFP or stdPCP–stdmScarlet. In the constructs, stem-loop cassettes were inserted in frame, preserving the open reading frame and avoiding frameshifts. The insertions did not interfere with splicing, as confirmed by RT–PCR, size verification by gel electrophoresis, and sequencing. **(D)** Representative images of transcription sites using a reporter with stem loops inserted downstream of the PTC within an exon. After overnight induction of transcription with PonA, time-lapse imaging was conducted for two hours. Each image shows an 8 × 8 µm² region centered on the transcription site. **(E)** Detection of transcription sites in cells expressing either the WWex or WPex construct, both containing stem loops inserted into the exon. Each dot represents the mean normalized intensity of transcription sites for wild-type (green) or PTC-containing (red) β-globin mRNA, measured over the indicated time period in 11 cells. Error bars indicate the standard deviation. Statistical analysis was performed using an unpaired, two-tailed t-test in GraphPad Prism.

Transcript retention may influence multiple stages of the RNA life cycle, including synthesis, processing, and nucleocytoplasmic distribution. To directly assess transcriptional output independently of RNA decay, we performed metabolic labeling of nascent RNA using the Click-iT system. Nascent transcripts were labeled for 30 min after 24 h of transcriptional induction, a time point at which NMD-mediated feedback is established. The short labeling period was chosen to minimize the contribution of NMD-dependent RNA degradation. Under these conditions, PTC-containing transcripts were produced at levels comparable to those of wild-type transcripts, suggesting that NMD feedback does not affect transcription initiation (Fig. 7A).

**Fig. 7.**
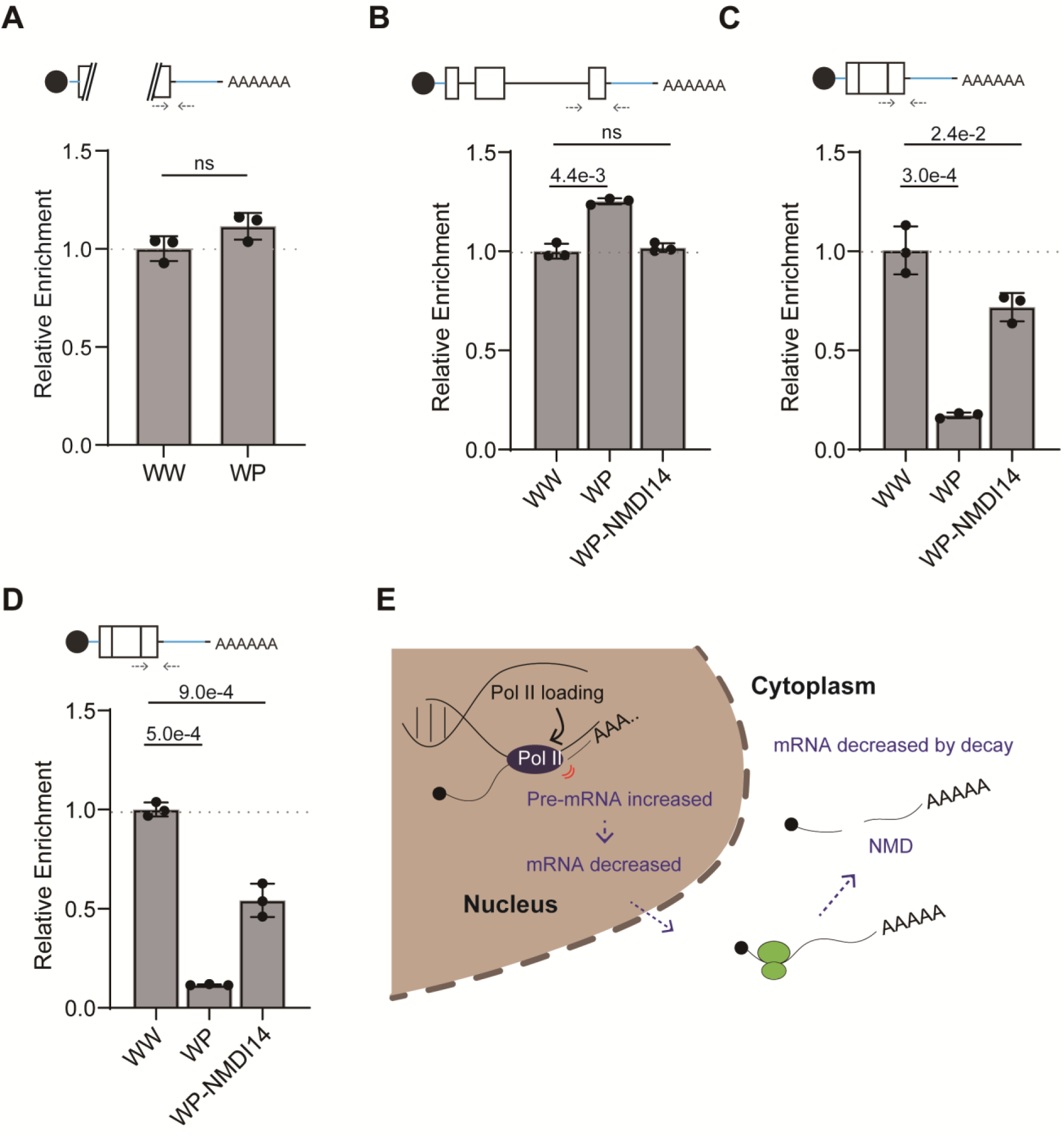
PTC-containing pre-mRNA and mRNA abundance changes without alteration of nascent transcript synthesis rate. **(A)** RNA synthesis rate is unchanged for PTC-containing globin–PP7 transcripts in WP compared with wild-type transcripts in WW. Nascent transcripts were captured for 30 min following 24 h of transcriptional induction to allow NMD feedback, using metabolic labeling with 5-ethynyl uridine (EU), a ribonucleotide analog containing an alkyne reactive group. The short labeling period was chosen to minimize detection of mRNA decay mediated by NMD. **(B)** Detection of PP7 pre-mRNA relative to MS2 pre-mRNA in WW, WP and WP treated with NMDI14 by RT–PCR in the nuclear fraction. **(C)** Detection of PP7 mRNA relative to MS2 mRNA in WW, WP, and WP treated with NMDI14 in the nuclear fraction. **(D)** Detection of PP7 mRNA relative to MS2 mRNA in the cytoplasmic fraction. Error bars indicate standard deviation (n=3). Statistical analyses were performed using unpaired, two-tailed *t*-tests in GraphPad Prism (ns, not significant). The positions of primer pairs are illustrated above each graph. **(E)** Schematic summary of changes in pre-mRNA and mRNA abundance induced by the presence of a PTC.

To examine how NMD-mediated feedback influences the subcellular distribution of transcripts, we measured pre-mRNA and mature mRNA levels in nuclear and cytoplasmic fractions in the presence or absence of the NMD inhibitor NMDI14 by RT-qPCR (Fig. 7B–D). The level of PTC-containing PP7 pre-mRNA was increased, consistent with previous reports, and this increase was reduced upon NMDI14 treatment, indicating that pre-mRNA accumulation is NMD-dependent (Fig. 7B).

In contrast, PTC-containing PP7 mRNA levels were lower in both nuclear and cytoplasmic fractions (Fig. 7C and D). The reduction of PTC-containing transcripts in the nuclear fraction is consistent with earlier studies (*35–39*). Taken together, these results suggest that NMD does not directly affect RNA synthesis but instead influences the nuclear pre-mRNA–to–mRNA balance (Fig. 7E).

To further dissect how this feedback influences transcription kinetics at the single-molecule level, we next analyzed the stochastic bursting behavior of the WT and PTC-containing alleles. Transcriptional activity was assessed by analyzing bursting dynamics (TS bursting), which reflect the timing and frequency of Pol II engagement at the transcription site. We quantified both the frequency of TS bursting (Fig. 8A and B) and the duration of ON-states (Fig. 8A and C). While the burst frequency was comparable between the wild-type and PTC-containing alleles, indicating that transcription initiation occurs at similar rates, the ON-state durations were significantly prolonged in the PTC-containing allele. To interpret this phenomenon, we developed a minimal stochastic model that captures key features of the observed transcriptional dynamics (Fig. S10). The model simulates ON-OFF transitions in fluorescence intensity using a set of state-dependent stochastic differential equations. Crucially, we introduced a mechanism for self-regulation in which bursts are temporally correlated, resulting in progressively longer ON durations. This extension recapitulates the experimental time series observed for the PTC-containing allele and provides a conceptual basis for understanding the altered dynamics. These results also suggest that the PTC mutation does not impact the frequency of Pol II recruitment but instead affects the kinetics of transcriptional progression but instead affects transcriptional progression and post-initiation processes that regulate transcript release from the transcription site. This interpretation is consistent with the fact that both WT and PTC transcripts are driven by the same bidirectional promoter. The increased ON-state duration may reflect delayed termination or Pol II release, potentially due to the retention of nascent RNA fragments produced by NMD. Our findings support a model in which the presence of a PTC triggers a feedback mechanism that prolongs the transcriptional engagement of Pol II, potentially reinforcing gene regulation at the level of elongation or transcript clearance. Based on this, we propose a model of transcriptional feedback triggered by NMD (Fig. 8D). Our data showed that transcriptional enlargement became apparent approximately one hour after the transcription site, which was first detected using the PonA-inducible system. Based on previously reported mRNA processing kinetics, the entire process from transcription to NMD-mediated decay is expected to occur as early as 30 minutes after transcription initiation. This suggests that the feedback mechanism can begin within that time window (Fig. 8E).

**Fig. 8.**
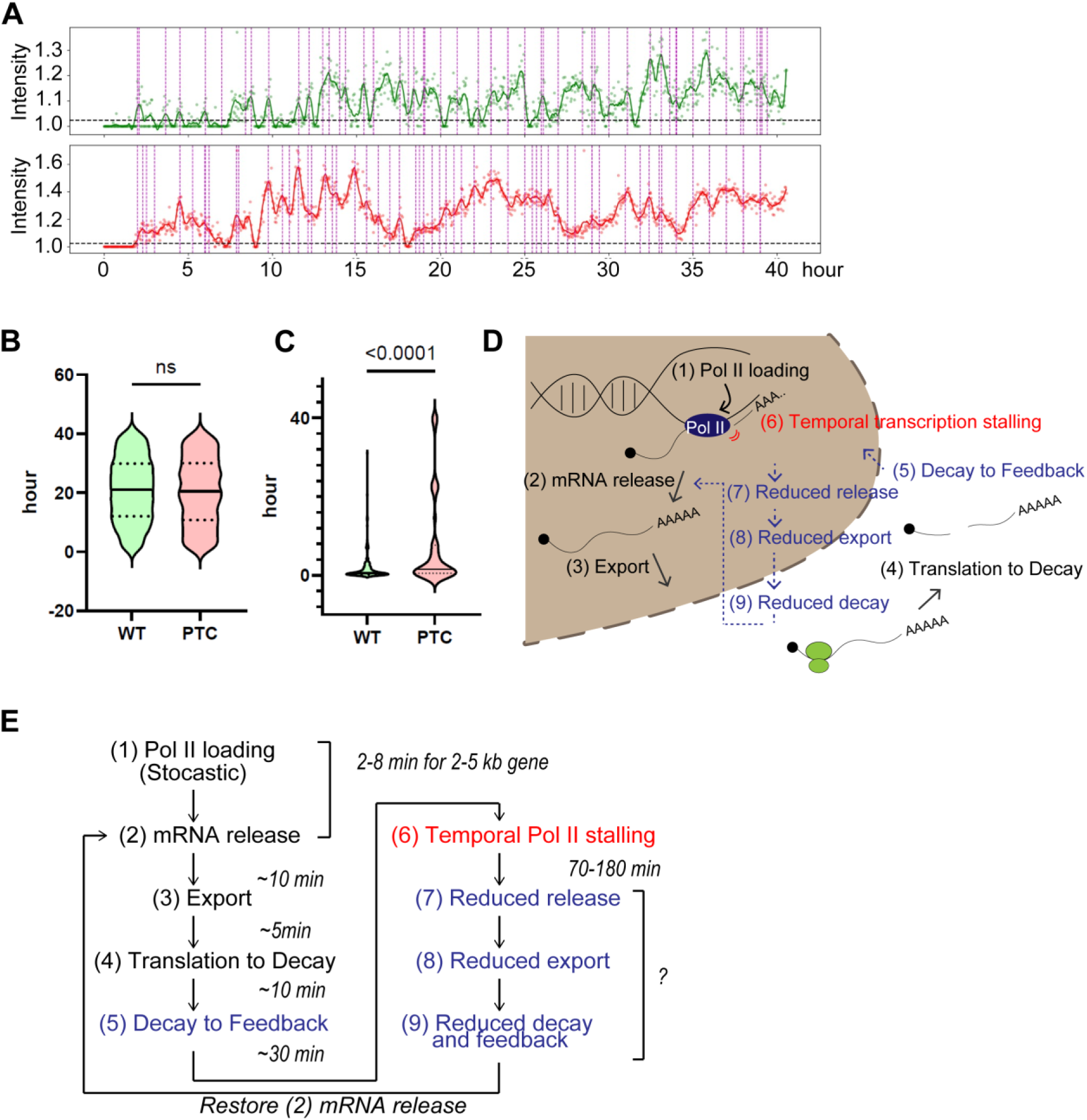
Model of feedback loop in NMD. (A) Example of transcription site (TS) intensity transition. Green and red lines represent the TS intensity of the wild-type and PTC-containing gene, respectively. Violin plot showing the frequency of transcriptional “on” state per hour **(B)** and duration of transcriptional bursting per hour **(C)**. Dotted lines represent quantiles; thick lines indicate medians. Statistical analysis was performed using an unpaired, two-tailed t-test (P<0.0001; ns indicates not significant). (D) Schematic representation of the stem-loop feedback mechanism in NMD. **(E)** Expected timing of transcriptional feedback. To estimate the expected timing, we evaluated the kinetics of mRNA from transcription to degradation. The transcription elongation rate ranges from 1.1 to 4.3 kb/min, with mRNA release typically completed within ∼5 min for a ∼4 kb transcript (*49*). Splicing can occur co-transcriptionally (*50*), the β-globin gene is relatively short, containing three exons and two introns. After transcription and maturation, mRNAs typically remain in the nucleus for ∼5 min before export. Export is rapid, with nuclear pore complex (NPC) transit times of ∼100–200 ms (*51*, *52*). Once in the cytoplasm, mRNAs are generally targeted for translation within 2–5 min (*53*). Although the timing of degradation by NMD varies depending on mRNA context, for a PTC located at codon 39 of β-globin mRNA, recognition of the PTC can occur within approximately ∼13 seconds (*54–56*). The entire process from transcription to translation and subsequent mRNA decay is expected to occur within ∼10 min (*45*, *46*)

We also observed that the transcriptional enlargement is transient, eventually diminishing over time (on average, within 1-2 hours, depending on the extent of the enlargement). This may be attributed to temporal stalling of Pol II, which would lead to reduced release of mRNA transcripts. As a result, downstream processes such as export, translation, decay, and subsequent feedback would also be diminished. Once the feedback weakens, the release of nascent mRNA likely returns to its normal rate.

## Discussion

Growing evidence suggests a strong link between RNA turnover and transcription (*1*, *42–44*). However, studying sequence-specific transcriptional feedback in NMD has been technically challenging due to the difficulty of linking nuclear with cytoplasmic events. In this study, we used simultaneous real-time imaging to detect transcriptional activities at both the wild-type and PTC-containing gene alleles, achieving high spatiotemporal resolution. Our findings provide robust evidence for rapid feedback from cytoplasmic decay to transcription in the nucleus. We also captured dynamic changes in the transcription site of the PTC-containing gene allele, driven by transcriptional stalling, which led to an increased spatial spread of nascent RNA at the transcription site. Importantly, this setup allows simultaneous monitoring of both alleles in the same cell, minimizing cellular variability and the challenges of comparing stochastic transcription bursting across different cells. A limitation of our system is that it relies on a reporter rather than an endogenous gene, and transcription is induced by a drug, which might not exactly mimic endogenous transcription regulation.

The phenomenon of transcriptional enlargement associated with NMD was first discovered using smFISH (*6*) to address a paradox: despite the requirement for translation, the reduction of PTC-containing mRNA was detected in isolated nuclear fractions (*35–39*). While it was later confirmed that NMD occurs on the cytoplasmic side, immediately after mRNA export (*45*), this discrepancy remains unresolved. Our study suggests that transcriptional stalling at the site of the PTC-containing gene may account for the reduction of PTC-containing mRNA in nuclear fractions, as it is likely due to pre-mRNA stalling rather than mRNA decay in the nucleus.

A related phenomenon is transcriptional adaptation (*7*, *8*, *25*, *26*), in which nonsense mutations lead to upregulation of homologous genes in response to cytoplasmic NMD. Although the underlying mechanisms remain unclear, this response has been observed across multiple systems and is thought to buffer genetic perturbations and maintain physiological function. Together, these phenomena share a common principle: NMD of cytoplasmic mRNA feeds back to influence transcription, linking cytoplasmic RNA surveillance to nuclear gene regulation.

One model proposed to explain transcriptional feedback in NMD involves the recognition of specific transcription sites by functional messenger ribonucleoprotein (mRNP) fragments that re-enter the nucleus. This possibility was previously explored by injecting in vitro-synthesized uncapped RNA, but the results were inconclusive (*7*, *8*). In our study, the use of a self-cleaving hammerhead ribozyme failed to trigger transcriptional enlargement, suggesting that RNA fragments alone are not sufficient to induce transcriptional enlargement. In contrast, transcriptional adaptation, defined as upregulation of homologous genes, has been reported when transcript cleavage is introduced by insertion of a T3H48 hammerhead ribozyme within an intron, as shown for *Actg2* or *UTRN* following cleavage of *Actg1* or *DMD* transcripts (*26*, *40*). Notably, the magnitude of transcriptional adaptation has been shown to vary across cell types. These studies further demonstrated that NMD is associated with increased chromatin accessibility, as detected by ATAC-seq, consistent with Pol II accumulation in an open chromatin environment. However, such chromatin opening was not detected at the PTC-containing locus when transcript cleavage was mediated by a hammerhead ribozyme (*40*). These observations raise the possibility that NMD-dependent Pol II stalling at nonsense alleles may involve mechanisms distinct from those operating during transcriptional adaptation, where the NMD target itself is transcribed, versus transcriptional adaptation, in which homologous genes are upregulated in trans. Further investigation will be required to identify the molecular components that mediate these distinct responses.

Additionally, the fact that nuclear import inhibition diminished the transcriptional enlargement of PTC-containing transcripts further supports the idea that a specific protein regulator may be involved in this mechanism, presumably by mediating the nuclear re-import of cytoplasmic RNA–protein complexes rather than RNA fragments alone.

Although RNA fragments generated by NMD are generally undetectable by biochemical methods such as northern blotting unless Xrn1 knockdown (*21*, *20*), this suggests that most intermediate fragments are rapidly and efficiently degraded. However, a previous study using live-cell imaging to analyze the decay kinetics of 3’fragment degradation by Xrn1 during NMD reported that Xrn1 occasionally dissociates from the RNA before completing degradation, implying that multiple rounds of Xrn1 engagement may be required for full transcript degradation (*46*). Such incomplete degradation could lead to the production of decay intermediate fragments, which may underlie a key mechanism by which these fragments regulate transcription. Notably, our results demonstrated that NMD-mediated transcriptional feedback is dependent on the nuclear import of proteins, indicating the involvement of protein factors in the mechanism. One potential candidate is Xrn1 itself, which has been implicated in linking RNA decay to transcriptional regulation in previous studies (*1*, *2*, *4*, *5*). Another is UPF1, which is known to associate with chromatin (*9*) and may play a role in this feedback loop. However, dissecting the specific contribution of these factors is challenging, as knockdown of Xrn1 or UPF1 not only disrupts the feedback mechanism but also impairs NMD activity itself. Therefore, further studies using refined experimental strategies will be required to identify the key protein factors that mediate NMD-coupled transcriptional feedback.

The biological significance of a feedback mechanism that couples cytoplasmic mRNA quality control with nuclear transcription remains to be fully elucidated. We propose that the transient stalling of Pol II observed in an NMD-dependent manner serves as a nuclear-cytoplasmic checkpoint. By delaying transcript release from the transcription site, this process could limit the export of mRNAs containing PTCs, thereby reducing the nuclear output of transcripts already destined for degradation and avoiding unnecessary metabolic costs. In this context, transient Pol II stalling may reflect the coordination of mRNA flux between the nucleus and the cytoplasm, ensuring that the rate of transcript release and export is aligned with the capacity of the cytoplasmic NMD machinery. This stalling may thus function as a reversible regulatory buffer that prevents the accumulation of aberrant mRNAs and preserves the efficiency of global mRNA processing.

While our study provides new insights into how NMD affects transcription dynamics, the precise nature of the feedback signal and the mechanisms underlying temporary Pol II stalling remain unclear. One possibility is that RNA molecules may have the potential to recognize specific genomic loci through RNA/DNA base pairing. One example of small RNA molecules regulating genomic DNA is the CRISPR-Cas system, which uses short (∼20 base pairs) RNA molecules to identify and cleave specific genomic loci. Additionally, small double-stranded RNAs, such as those involved in RNA interference (RNAi) pathways (e.g., small interfering RNAs (siRNAs), PIWI-interacting RNA (piRNA)), are known to regulate chromatin modifications (*47*, *48*), suggesting a common mechanism for small RNAs in chromatin regulation. Further studies will be necessary to elucidate the molecular details of these processes and to determine whether similar mechanisms contribute to the feedback observed in NMD-mediated regulation.

## Materials and Methods

### Cell lines and tissue culture

The human U2OS FRT cell line was established in a previous study (*30*). U2OS FRT PonA cell line was generated by stable transfection with the pERV3 plasmid, which expresses the synthetic VP16-glucocorticoid/ecdysone receptor (VgEcR) and retinoid X receptor (RXR), both required for activation of transcription from the PonA promoter, as previously described (*29*). Bidirectional PonA reporters constructs pFRT-PonA-BI-Gl-WT-24xMS2-WT-18xPP7 (WW) and pFRT-PonA-BI-Gl-WT-24xMS2-PTC-18xPP7 (WP) were described previously (*29*) and integrated into a single defined genomic locus using the Flp-In system (Thermo Fisher Scientific). Correct splicing of WW and WP transcripts in stably expressing U2OS FRT PonA cell lines was verified by RT–PCR followed by gel electrophoresis and Sanger sequencing. Proper 3’ end processing was confirmed by 3’ RACE (Rapid Amplification of cDNA Ends (TaKaRa, Cat# TKR-6121). Additional constructs were generated in this study, including pFRT-PonA-BI-Gl-WT-24xMS2-HHR-18xPP7 (WH), pFRT-PonA-BI-mu-WT-24xMS2-WT-18xPP7 (mu-WW), pFRT-PonA-BI-mu-WT-24xMS2-PTC-18xPP7 (mu-WP), pFRT-PonA-BI-Gl-WT-6xMS2exon-WT-5xPP7exon (WWex), and pFRT-PonA-BI-Gl-WT-6xMS2exon-PTC-5xPP7exon (WPex). For the WH construct, a synthetic hammerhead ribozyme sequence (gacgagcttactcgtttcgtcctcacggactcatcag) was inserted upstream of the PP7 stem-loop cassette. For the mu-WW and mu-WP constructs, the mini-μ gene was obtained from a previous study (*9*) and cloned under the same bidirectional PonA promoter. For the WWex and WPex constructs, the sequence downstream of the PTC in the WW and WP reporters was replaced with the endogenous human β-globin 3’UTR. Subsequently, 6xMS2 and 5xPP7 stem-loop cassettes were inserted in frame within exon 1 to preserve the open reading frame. Correct splicing was verified by RT–PCR, size analysis by gel electrophoresis, and Sanger sequencing. Stable cell lines were generated by co-transfection with pOG44 followed by hygromycin selection. All cell lines were maintained at 37 °C with 5% CO₂ in Dulbecco’s Modified Eagle Medium (DMEM) supplemented with 4.5 g/L glucose, 10% fetal bovine serum (FBS), and 1% penicillin-streptomycin.

### Live cell imaging acquisition

For live-cell imaging, the culture medium was replaced with L-15 medium supplemented with 10% fetal bovine serum (FBS) and 1% penicillin-streptomycin prior to imaging. Wide-field fluorescence images were acquired using either an IX-81 inverted microscope (Olympus), equipped as previously described (*50*), or an ECLIPSE Ti2-E inverted microscope (Nikon) equipped with a four-laser unit (LUD-H4) and an ORCA-Fusion BT CMOS camera (Hamamatsu). The Olympus system was operated with MetaMorph software using a 60×/1.4 NA oil immersion objective, while the Nikon system was controlled by NIS-Elements software and used a CFI Apo TIRF 60XC Oil objective (MRD01691). During imaging, cells were maintained at 37 °C using a stage-top incubator (INUBH-ZILCS-F1, Tokai Hit, Japan).

Optical sectioning was performed using a 500 nm Z-step over a total depth of 5.0 µm. The exposure time was set to 50 milliseconds for each optical plane and channel. To inhibit translation, nonsense-mediated mRNA decay (NMD), or nuclear import of proteins, cells were treated with cycloheximide (CHX, Tocris Bioscience, Cat# 0970) at 100 µg/ml, NMDI14 (Sigma-Aldrich, Cat# SML1538) at 40 µM, or importazole (Sigma-Aldrich, Cat# 401105) at 50 µM, according to the treatment timelines described in the corresponding figure legends.

For live-cell widefield imaging, laser power was set to 1 mW (488 nm) and 0.9 mW (561 nm) when measured before the objective lens. After accounting for internal optical losses, the estimated laser power at the sample plane was on the order of several tens of microwatts.

To determine the spatial distance between transcription sites, 3D volumes were acquired using an Andor Dragonfly spinning-disk confocal microscope equipped with a Nikon CFI Apochromat TIRF 100x Oil objective (NA 1.49). Z-stacks were captured over an 8 µm range with a 0.2 µm axial step size. Following acquisition, the 3D data were deconvolved and imported into Imaris software (Oxford Instruments), where transcription sites were manually identified. Inter-site distances were calculated using the 3D coordinates of the signal centroids, specifically targeting the voxel of peak fluorescence intensity for each spot. For confocal imaging, laser power was measured directly after the objective lens using a Thorlabs PM100D power meter, with average power levels maintained at 25 mW (488 nm) and 30 mW (561 nm). Due to the high-speed scanning of the spinning-disk system, the sample was subjected to pulsed illumination.“

### UPF1 knockdown

UPF1 knockdown was performed using UPF1-targeting shRNA and control shRNA, as described previously (*31*). U2OS FRT PonA WW cells were transfected using a laboratory-prepared PEI-based transfection reagent (referred to as PEI-max). Cells were seeded at approximately 70-80% confluency in complete growth medium one day prior to transfection. For each well of a 6-well plate, plasmid DNA was diluted in 100 µL of Opti-MEM (Thermo Fisher Scientific), and 6 µL of PEX-max solution was added. The DNA and reagent mixture was incubated at room temperature for 15 min to allow complex formation, then added dropwise to the cells. Transfected cells were maintained at 37 °C with 5% CO₂ and harvested 48 hours post-transfection for analysis.

### Chromatin immunoprecipitation (ChlP)

ChIP was performed using the EpiQuik™ Chromatin Immunoprecipitation Kit (catalog no. P-2002; Epigentek) according to the manufacturer’s instructions. Briefly, 3×10^6^ U2OS_FRT_PonA stably expressing WW or WP cell lines were crosslinked with 1% formaldehyde containing cell culture medium and lysed in presence of protease inhibitor cocktail. Chromatin pellet was resuspended in 200 µL of resuspend buffer and incubate on ice for 10 min following by occasional vertexing. Chromatin fragmentation was followed by sonication (60 cycles of 30 s on and 30 s off) to an average DNA size of 200-600 bp, as confirmed by 2% agarose gel. An aliquot (10 µL) of fragmented chromatin was reserved as input DNA. The remaining chromatin (190 µL) was subjected to immunoprecipitation using an anti-RNA polymerase II antibody (RPB1; catalog no. 664906; BioLegend), with non-immune IgG used as a negative control separately. Following extensive washing, chromatin complexes were eluted after reverse-crosslinking at 65 °C for 90 min with proteinase K treatment. DNA was purified using the kit-provided spin columns and eluted in 20 µL of nuclease-free kit buffer for downstream quantitative PCR analysis. Quantitative PCR was performed using PowerTrack™ SYBR Green Master Mix (catalog no. A45109; Thermo Fisher Scientific) according to the manufacturer’s instructions with 10 µL reaction scale.

### Subcellular fraction of cytoplasmic and nuclear RNA

Cell fractionation was performed on ice or at 4 °C in the presence of RNase inhibitor (code no. 315-08121) using cell pellets corresponding to 2 × 10⁶ cells. Pellets were gently resuspended in 100 µl cytoplasmic lysis buffer (0.15% NP-40, 10 mM Tris-HCl, pH 7.0, 150 mM NaCl) and incubated on ice for 5 min. Lysates were layered onto 300 µl sucrose buffer (10 mM Tris-HCl, pH 7.0, 150 mM NaCl, 25% sucrose) and centrifuged at 16,000g for 10 min. The supernatant corresponding to the cytoplasmic fraction was carefully collected and mixed with an equal volume of TRI Reagent (TRI Reagent®, catalog no. TR 118).

Nuclei were rinsed in 200 µl nuclear wash buffer (0.1% Triton X-100, 1 mM EDTA in 1×PBS) and collected by centrifugation at 1,500g for 1 min. The resulting pellet was resuspended in 100 µl glycerol buffer (20 mM Tris-HCl, pH 8.0, 75 mM NaCl, 0.5 mM EDTA, 50% glycerol, 0.85 mM DTT) and combined with 250 µl TRI Reagent. Cytoplasmic and nuclear homogenates were incubated at room temperature for 5 min, followed by vigorous shaking for 15 s after the addition of 100 µl chloroform. Samples were incubated at room temperature for 10 min and centrifuged at 12,000g for 15 min at 4 °C. The aqueous phase was transferred, mixed with an equal volume of isopropanol, and incubated at room temperature for 5 min. RNA was pelleted by centrifugation at 12,000g for 15 min at 4 °C, washed with 75% ethanol, and resuspended in RNase-free water. RNA concentration was measured using a NanoDrop One spectrophotometer (Thermo Fisher Scientific) and subsequently used for cDNA synthesis.

### Nascent RNA labeling and capture

Nascent RNA was labeled and isolated using the Click-iT Nascent RNA Capture Kit (Thermo Fisher Scientific, catalog no. MP10365). U2OS FRT PonA cells stably expressing WW and WP reporter constructs were cultured under standard conditions and induced with ponasterone A (ponA, 10 µM) for 24 h, followed by labeling with 0.5 mM 5-ethynyl uridine (EU) for 30 min to label newly synthesized RNA. Following labeling, cells were harvested, and total RNA was extracted according to the manufacturer’s protocol. EU-labeled RNA was conjugated to biotin azide via copper(I)-catalyzed click chemistry and selectively captured using streptavidin magnetic beads. After stringent washing, nascent RNA was eluted and used directly for downstream reverse transcription and quantitative analysis.

### Real-time RT-qPCR

Total RNA was purified using either TRIzol™ Reagent (Thermo Fisher Scientific, Cat# 15596018) or TRI Reagent (Cosmo Bio, Cat# TR118), following the manufacturers’ instructions. For reverse transcription PCR (RT-PCR), RNA was treated with RNase-free DNase I (1 U/μL; Thermo Fisher Scientific, Cat# EN0521) for 30 min at 37 °C. cDNA was synthesized from 0.2 µg of total RNA in a 25 µL reverse transcription reaction using SuperScript III Reverse Transcriptase (Invitrogen) with random hexamers, or using the GeneAce cDNA Synthesis Kit (Nippon Gene, Cat# 319-08881), according to the respective manufacturer’s protocol.

Quantitative PCR (qPCR) was performed using either PowerUp SYBR Green Master Mix (Thermo Fisher Scientific, Cat# A25741) or PowerTrack™ SYBR Green Master Mix (Thermo Fisher Scientific, Cat# A45109). Relative gene expression levels were calculated using the ΔΔCt method, with beta-actin used as the housekeeping gene for normalization of UPF1 knockdown. Data were representative of at least three independent biological replicates. Primer sequences are provided in the supplementary table.

## Data analysis

Time-lapse images of transcription were obtained by generating maximum intensity projections from the z-series. Images were acquired at 2-minute intervals, with total imaging duration varying by experiment as indicated in the corresponding figure legends. Transcription sites were detected and tracked using Trackmate (*32*). with the Simple LAP tracker. Fluorescence intensities of the transcription sites were measured and normalized to the diffusive EGFP or mScarlet signal in the nucleus. The normalized spot intensities at each time point were plotted and smoothed using R. Transcription burst peaks were identified using the *findpeaks* function in R.

The statistics of burst events, such as their duration, frequency, the intensity time-series were gathered after further analysis as follows. The signal (intensity time-series), sans the outliers, was smoothened first using a Savitzky-Golay filter. For both cases, with and without PTC, a polynomial of order 3 with a time-window around 40 frames produced reasonable smoothing. Defining an ON-state above a certain Intensity threshold, its duration and related statistics were computed. The bursts were identified as the local maxima of the original signals. The closely spaced bursts (within three consecutive frames) are then replaced by their means to reduce overestimation. The bursts identified, the statistics of burst separation, and burst frequency per hour are computed for all observed signals.

### Simulation of self-regulated stochastic process

To model the observed transcriptional dynamics, we developed a simplified stochastic framework that captures the essential features of fluorescence intensity fluctuations seen in our experiments. In both wild-type (WT) and premature termination codon (PTC)-containing alleles, the fluorescence intensity exhibits stochastic transitions between an OFF state (characterized by a baseline intensity) and an ON state (characterized by elevated intensity). These ON and OFF states occur with randomly distributed durations. Notably, transcription from the PTC-containing allele displays significantly longer ON durations compared to WT.

Based on prior knowledge of transcriptional bursting, we describe the fluorescence intensity time series *I(t)* using the following stochastic differential equation:

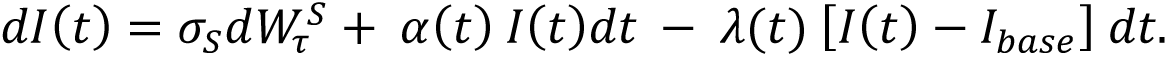

Here, *I(t)* denote the observed fluorescence intensity. The first term represents a Wiener process dWτ^S^, where S∈{ON, OFF} and σ_S_ denotes the noise amplitude associated with each state. The second term, active only during the ON state, represents an intensity growth process governed by a time-dependent rate function α(t). The third term accounts for intensity decay toward a baseline value *I*_base_, with a decay rate λ(t). During the OFF state, both α(t) and λ(t) are set to zero.

Although this stochastic model is a phenomenological approximation, it serves as a useful proxy for the underlying biochemical network, which is currently too complex to resolve using our experimental setup. However, our data suggests that this network may introduce correlations in ON-state duration, particularly evident in the transcriptional behavior of the PTC-containing allele.

To incorporate such correlations, we introduce a surrogate equation that governs the evolution of the ON-state duration τ_ON_, conditioned on the change in fluorescence intensity:

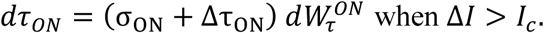

Here Δ*I* = *I*(*t* + 1) − *I*(*t*), and *I*_*c*_ is a threshold value set arbitrarily. When Δ*I*>*Ic*, the model introduces a small random increment Δτ_ON_, favoring prolonged ON durations in response to large intensity changes. This condition is based on the intuition that in a Wiener process, large changes between consecutive steps are rare and thus may encode weak correlations. Accordingly, 1/*I*_*c*_ can be interpreted as a proxy for the strength of this correlation. This minimal stochastic description successfully recapitulates the key features of the transcriptional dynamics observed in our experimental data. Importantly, it provides a conceptual basis for understanding the mechanistic differences between WT and PTC-containing transcriptional activity.

## Acknowledgments

We thank members of the Singer laboratories for discussions, the Einstein FACS and Genomics cores. This work was begun at Einstein supported by 5R35GM136296 to RHS. This work was also supported by the World Premier International Research Center Initiative (WPI), MEXT, Japan; the Naito Grant for Female Scientists; the Mitani Foundation for Research Grant to HS; and the Japan Society for the Promotion of Science (JSPS KAKENHI Grant Number 24K23179 to TD).

## Author contributions

Conceptualization: HS, RHS

Methodology: HS, TD

Investigation: MDI, HS, TD

Formal analysis: MDI, TD, HS

Visualization: HS, TD

Funding acquisition: RHS, HS

Project administration: RHS, HS

Resources: HS, RHS

Supervision: RHS, HS

Writing – original draft: HS

Writing – review & editing: MDI, RHS, HS, TD

## Data and materials availability

All data and code necessary to evaluate and reproduce the results reported in this paper are available in the manuscript and the supplementary materials. Plasmids, engineered cell lines, and other materials generated during this study are available from the corresponding author upon reasonable request.

## Supplementary Materials

Figs. S1 to S10

Supplemental table: primer list for RT-qPCR

Captions for Movies S1 to S2

Movies S1 to S2 (separate file)

