## Supplementary_Materials for "Real-time imaging of transcriptional feedback in nonsense-mediated mRNA decay"

**This PDF file includes:**

Figs. S1 to S10
Supplemental table: primer list for RT-qPCR
Captions for Movies S1 to S2

**Other Supplementary Materials for this manuscript include the following:**

Movie S1: area5923_Movie Still_mov1_seq1_v1.avi
Movie S2: area5923_Movie Still_ seq2_v1.avi
All data sets in this study

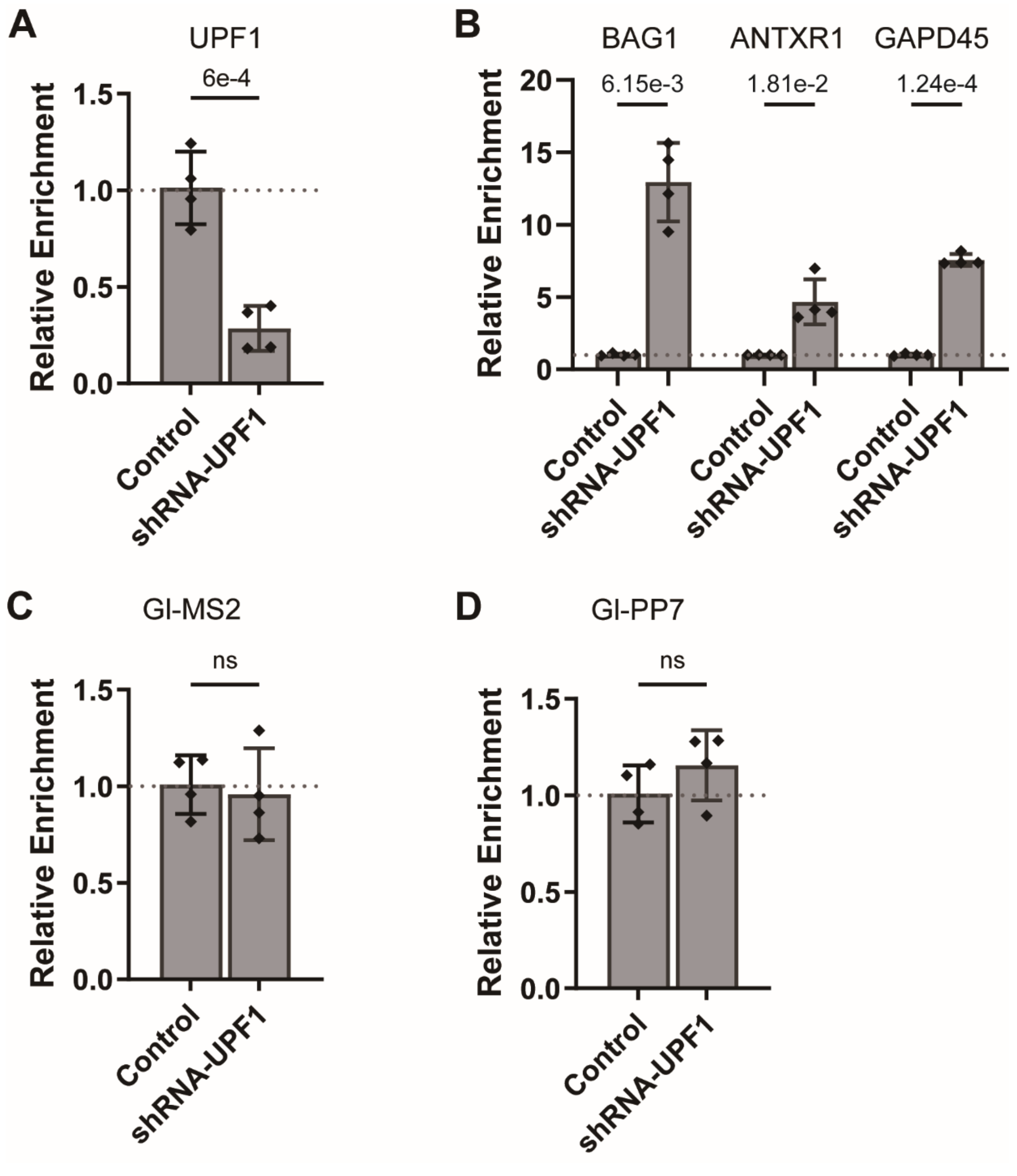


**Fig. S1. Insertion of multiple stem loops into the 3' UTR does not induce NMD in the Gl reporter.** (A) Confirmation of shRNA-mediated UPF1 knockdown by RT-qPCR. β-actin mRNA served as an internal reference. (B) Quantitative detection of endogenous NMD targets following UPF1 downregulation. (C and D) RT-qPCR quantification of β-globin mRNA containing (C) MS2 stem-loops or (D) PP7 stem-loops in WW-expressing cells. β-actin mRNA was used as a normalization control. P-values were determined by two-tailed unpaired Student's t-tests (ns, not significant). Error bars represent the standard deviation (SD) from at least three independent experiments. Statistical analyses were performed using GraphPad Prism software.

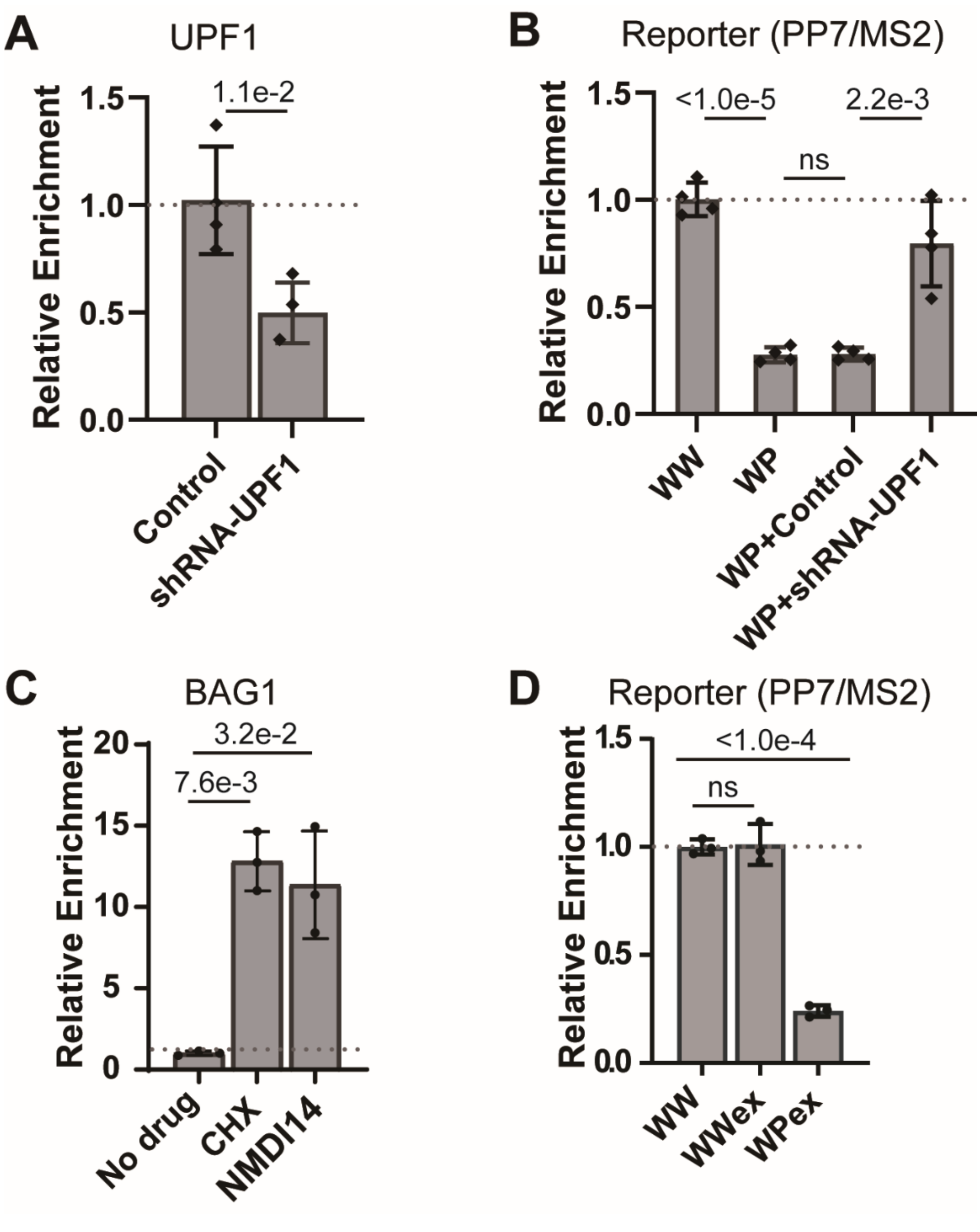


**Fig. S2. PTC-containing Gl-PP7 transcripts are NMD sensitive.** (A) Confirmation of shRNA-mediated UPF1 knockdown by RT-qPCR. β-actin mRNA served as an internal reference. (B) Quantitative detection of Gl-PP7 transcripts following UPF1 downregulation. (C) Quantitative detection of endogenous NMD targets following CHX and NMDI14 treatment. (D) Quantitative measurement of Gl-PP7 transcripts following expression of the WWex and WPex constructs. P-values were determined by two-tailed unpaired Student's t-tests (ns, not significant). Error bars represent the standard deviation (SD) from at least three independent experiments. Statistical analyses were performed using GraphPad Prism software.

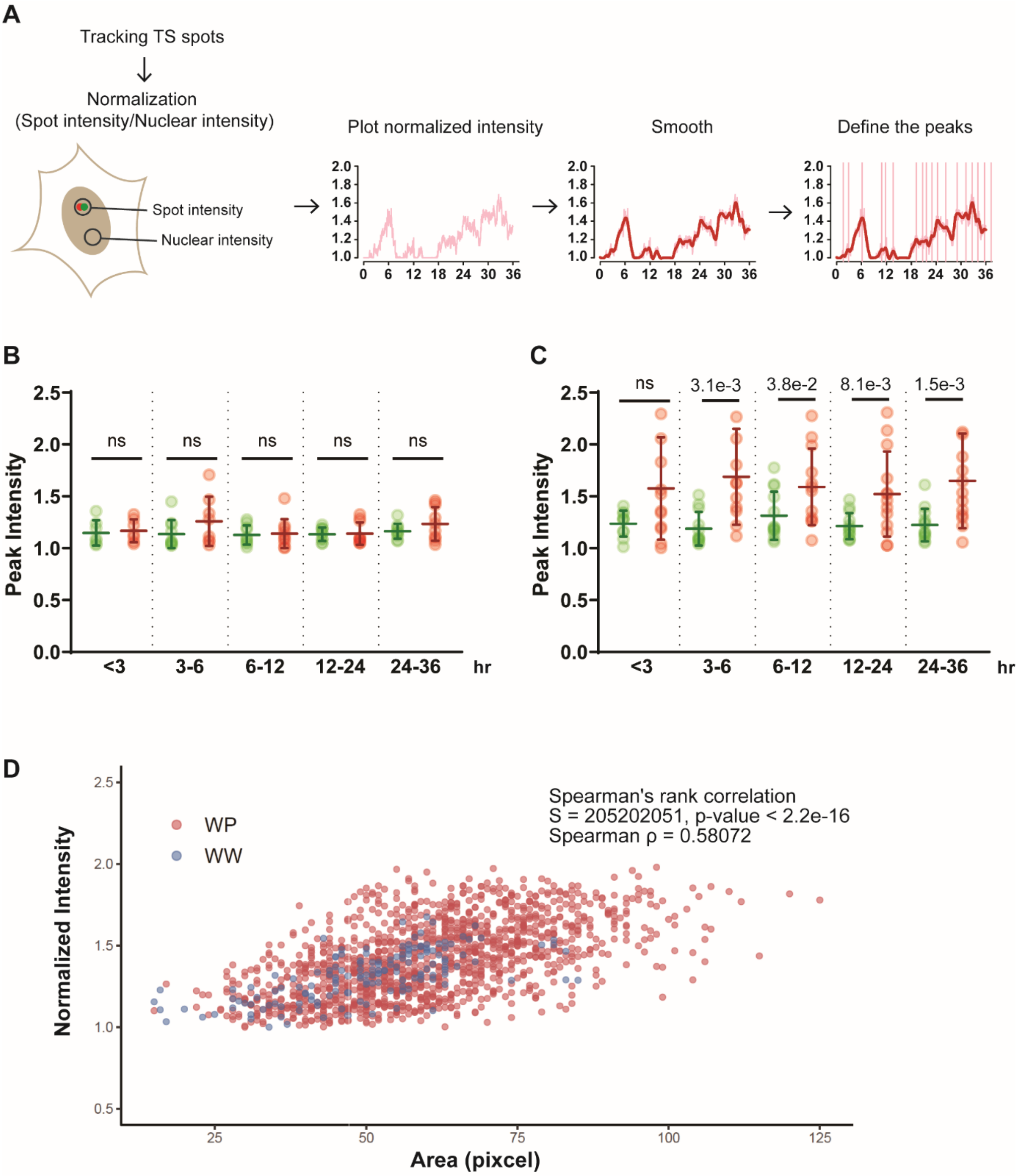


**Fig. S3. Analytical scheme of peak detection and peak analysis of transcription sites. (A)** Schematic of analytical scheme for transcriptional activity. Transcription sites bidirectionally expressing mRNA labeled with MS2 or PP7 system were imaged every 2-min for more than 36-hours in live cells and detected using TrackMate spot analysis software including the extension of TrackMate-extras for multi-channel tracking. Transcription sites labeled with green or red fluorescent proteins were tracked in the same ROI between each color except when transcription sites of each color expressing bi-directionally did not overlap. When one of the transcription sites expressing bidirectional was not detectable, a transcription site labeled with another color was used to define the ROI. When both transcription sites were temporally invisible, the action of “Close gaps by introducing new spots”, which introduces the new spots based on the positions and size calculated using the linear interpolation from the track, was used. The mean intensity of identified spot at each time was normalized by dividing the spot intensity by the nuclear intensity. The normalized intensities of transcription sites were plotted, smoothed using loess regression and smoothing in R. The peak (transcription burst) was defined using the findpeaks function in R (pracma: Practical Numerical Math Functions) (*57*). The code using this analysis is available upon request. Detection of transcription sites expressing WW (B) or WP (C) construct for 36-hours. Single dots denote the mean of normalized intensity of transcription sites during indicated time duration from single cells. P-values were determined using two-tailed unpaired t-tests (ns, not significant) from 10 (WW) or 13 (WP) cells. Error bars = Standard deviation in cell populations. (D) Relationship between transcription site (TS) area and TrackMate-quantified fluorescence intensity. Scatter plots show TS area (pixels) versus normalized TS fluorescence intensity for WW and WP constructs.

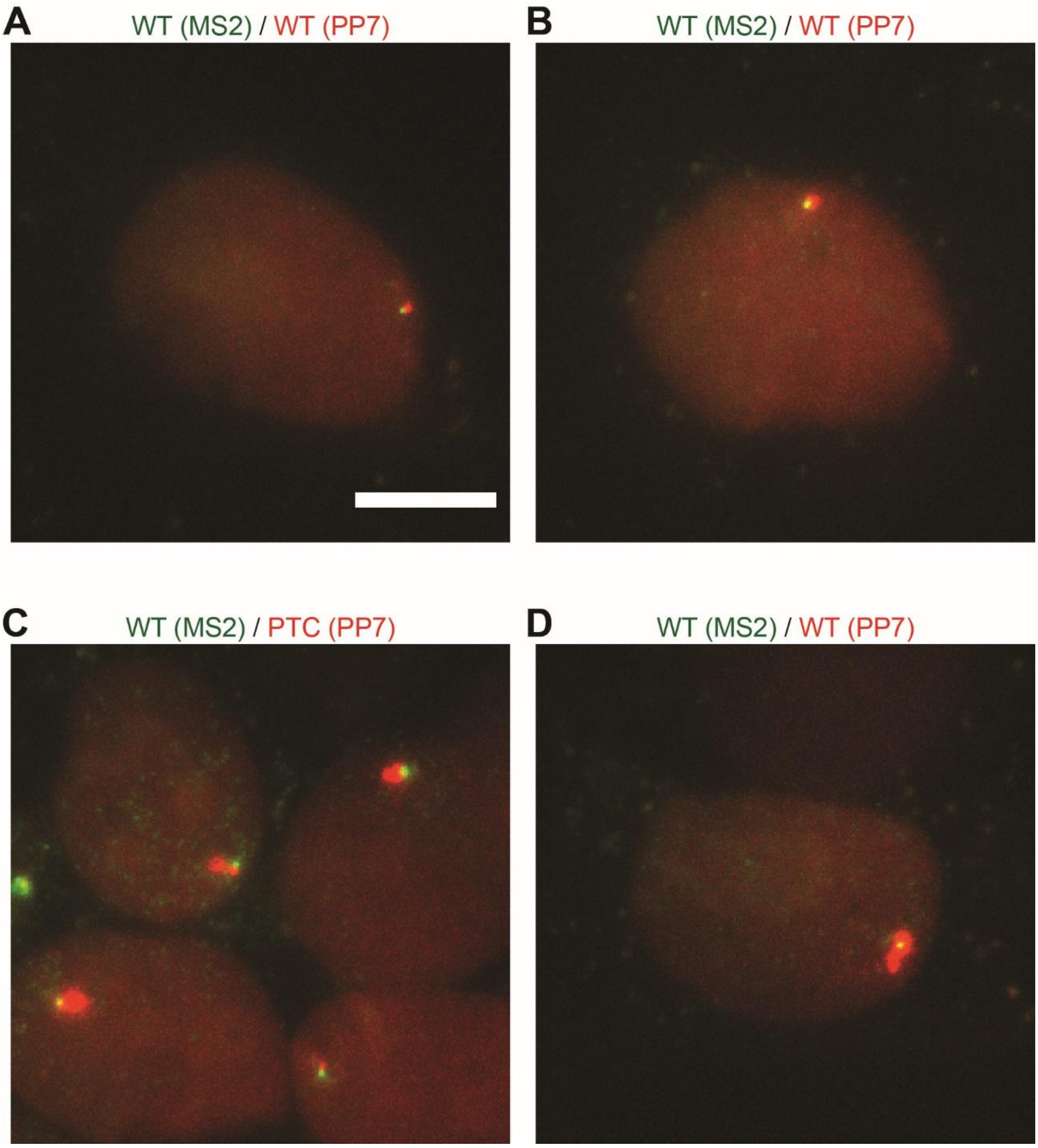


**Fig. S4. Simultaneous detection of transcription sites at the Igμ NMD reporter with and without PTC.** (A–B) Transcription sites expressing wild-type Igμ mini reporter mRNA labeled with MS2 in the 3'UTR (green) and PP7 in the 3'UTR (red) from both directions of the bidirectional promoter. (C–D) Transcription sites expressing either wild-type Igμ with MS2 in the 3' UTR (green) or PTC-containing Igμ with PP7 in the 3'UTR (red) from each direction of the promoter. Scale bar = 10 μm.

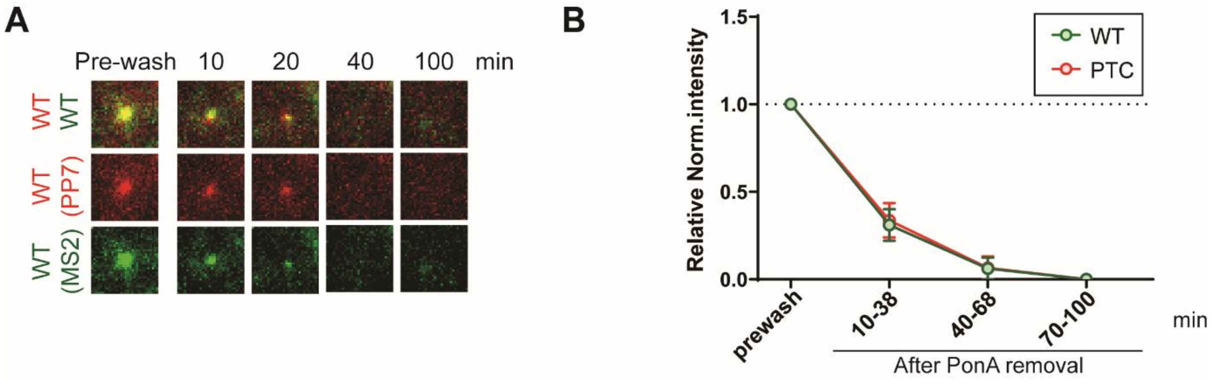


**Fig. S5. Rapid disappearance of transcription sites in WW cells following PonA removal.** (A) Representative images of transcription sites before (Pre-wash) and after PonA removal (10–100 min). Each image displays a 7 × 7 μm² field of view. The experimental timeline follows the protocol described in Figure 3. (B) Quantification of transcriptional activity from the WP construct before (Pre-wash) and after PonA removal (10-100 min). Each data point represents the mean normalized intensity of transcription sites across 11 cells. Error bars represent the standard deviation (SD) of the cell population. Statistical analysis was performed using an unpaired, two-tailed t-test in GraphPad Prism. P-values for the 11–38 min, 40–68 min, and 70–100 min intervals were all non-significant.

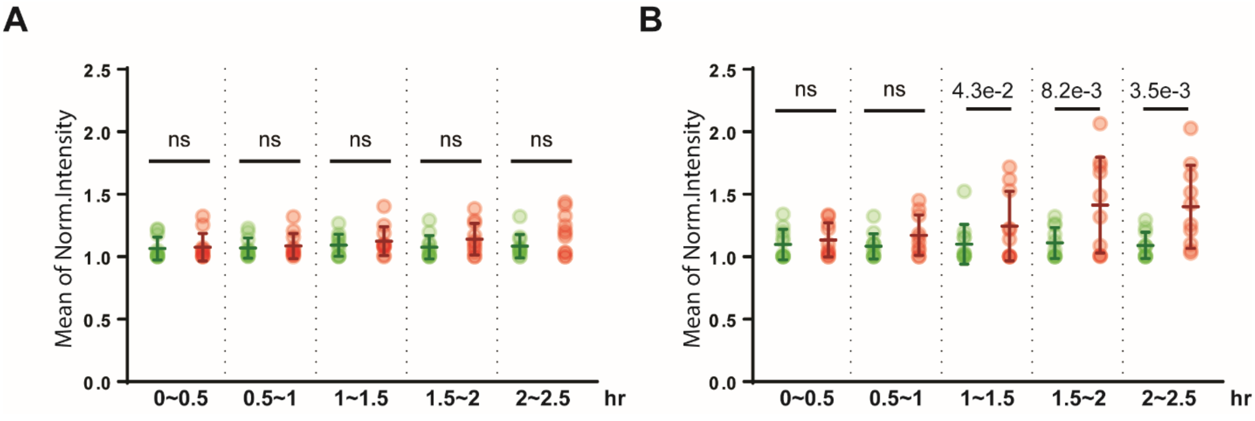


**Fig. S6. Transcriptional enlargement begins about 1 hour after transcription was initiated.** Detection of transcription sites expressing WW (A) or WP (B) construct. Single dots denote the mean of normalized intensity of transcription sites during indicated time duration from single cells. P-values were determined using two-tailed unpaired t-tests (ns, not significant) from 11 (WW) or 10 (WP) cells. Error bars = Standard deviation in cell populations.

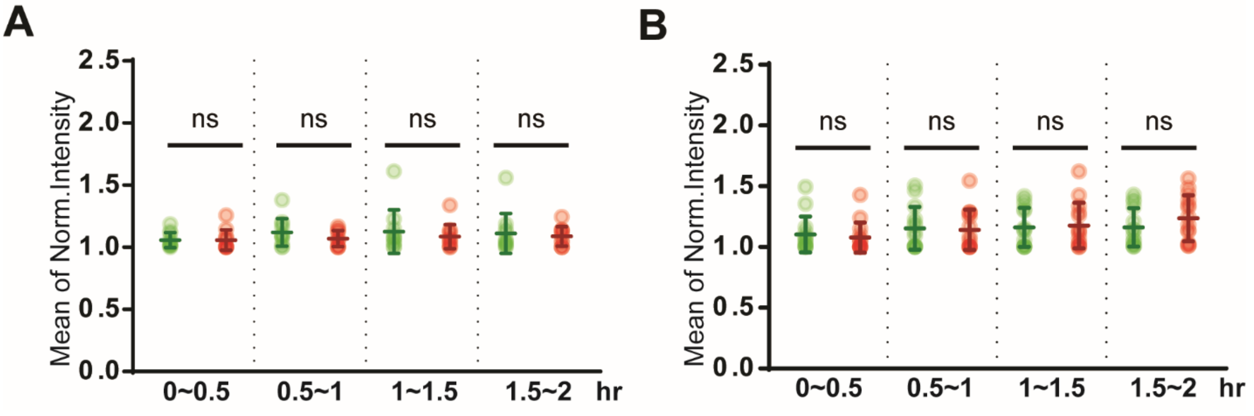


**Fig. S7. Detection of transcription sites expressing the WW or WP construct with the translation inhibitor cycloheximide (CHX).** Detection of transcription sites expressing WW (A) and WP (B) construct with CHX. Single dots represent the mean normalized intensity of transcription sites from individual cells (green: wild-type; red: PTC-containing β-globin transcription sites) during the indicated time intervals. P-values were determined using two-tailed unpaired t-tests (ns, not significant,). Error bars indicate standard deviation across cell populations. n = 14.

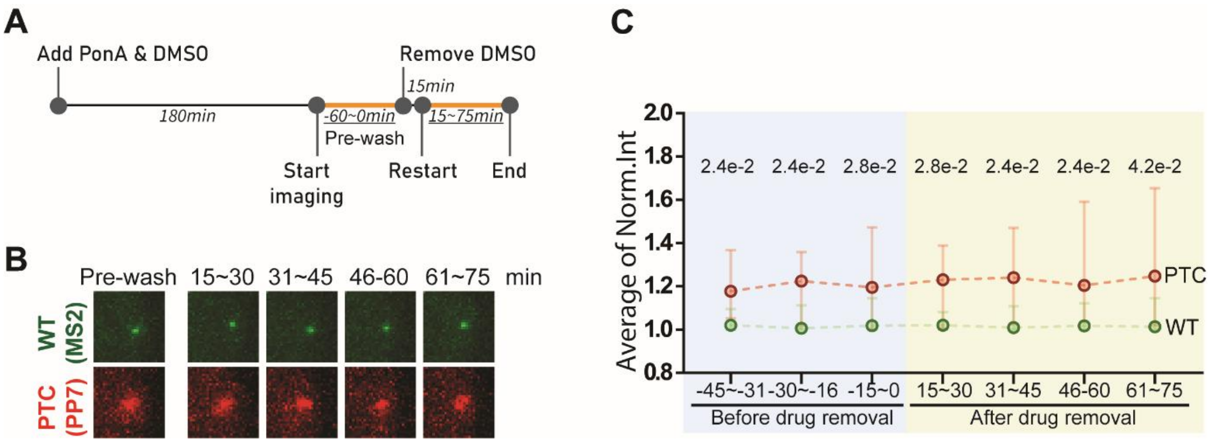


**Fig. S8. Changing medium does not trigger PTC-specific transcriptional enlargement.** (A) The timeline of transcription induction by supplement of PonA with DMSO and following DMSO removal. Orange lines and underlined times indicate the time duration for real-time imaging. (B) Images of transcription sites before (Pre-wash) and after removal of DMSO (15-75 min). The image size of each transcription site shown here is 8 x 8 μm$^2$. (C) The detection of transcription activities expressing WP construct before (Pre-wash) and after removal of DMSO (10-90 min). Single points with 95% confidence intervals indicate the median normalized intensity for wild-type (green) and PTC-containing β-globin transcription sites (red) across 13 cells at each indicated time window. Statistical analysis was performed in GraphPad Prism using paired, nonparametric Wilcoxon matched-pairs signed-rank tests (ns, not significant).

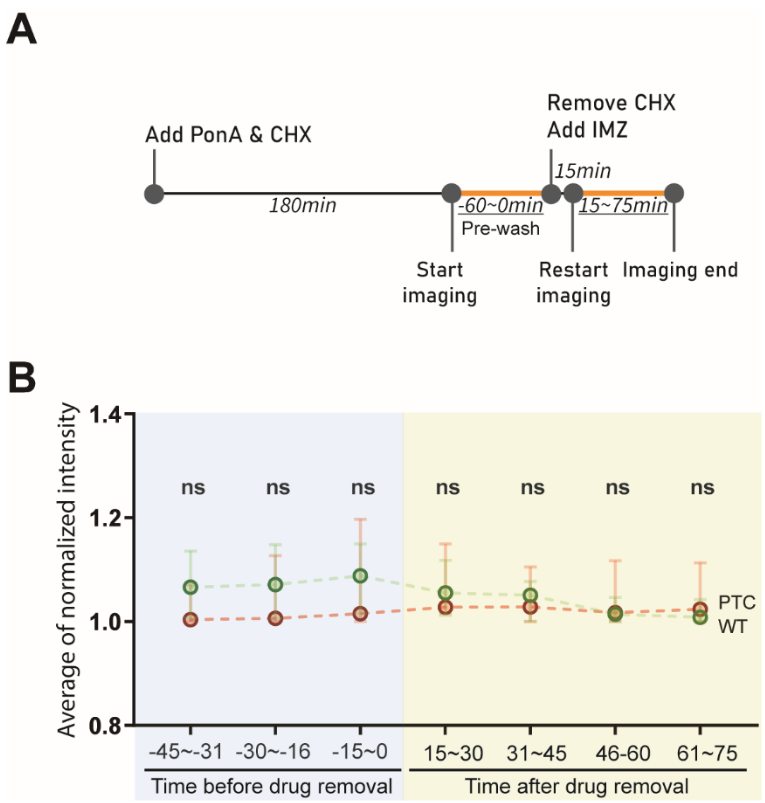


**Fig. S9. Nuclear transport importin-β inhibitor, importazole inhibited PTC-specific transcription enlargement.** (A) The timeline of transcription induction by supplement of PonA with translation inhibitor, CHX and following the removal of CHX and supplement of importin β inhibitor, 50 μM importazole (IMZ) (*42*). (B) The detection of transcription sites expressing WP construct before and after removal of CHX with the supplement of IMZ. Single points with 95% confidence intervals indicate the median normalized intensity for wild-type (green) and PTC-containing β-globin transcription sites (red) across 12 cells at each indicated time window. Statistical analysis was performed in GraphPad Prism using paired, nonparametric Wilcoxon matched-pairs signed-rank tests (ns, not significant).

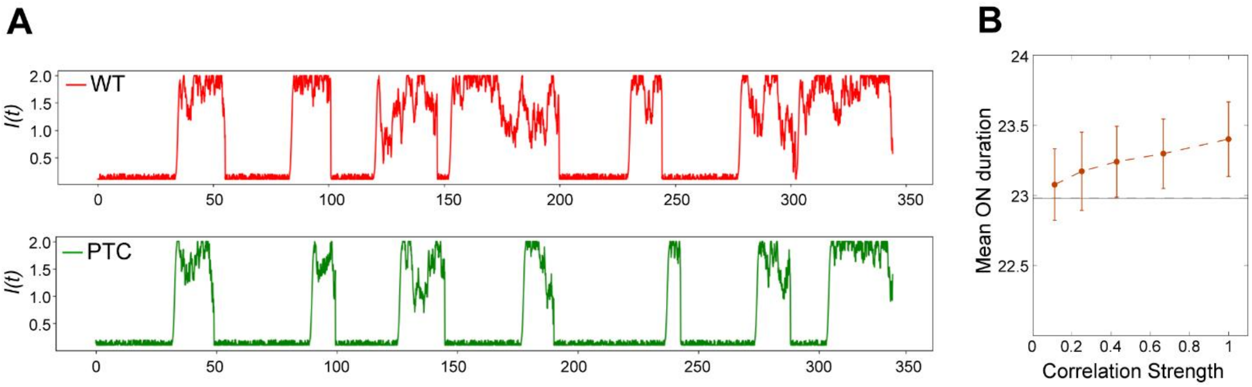


**Fig. S10. Transcription as a self-regulated stochastic process.** (A) Simulated time series of fluorescence intensity $\boldsymbol{I(t)}$ illustrating a single realization of the stochastic ON-OFF transcriptional process. The red trace represents the unregulated model, capturing the burst dynamics characteristic of the wild-type (WT) allele. The green trace demonstrates the effect of incorporating self-regulation via correlated burst durations, resulting in extended ON-state durations that resemble the transcriptional behavior observed for the PTC-containing allele. (B) The ON-state duration increases consistently with the strength of correlation, supporting the hypothesis that enhanced burst correlation underlies the prolonged transcriptional activity seen in the PTC context. This provides a conceptual explanation for the dynamic differences between WT and PTC-containing alleles, as discussed in the main text.

**Supplemental table**

Primer list for RT-qPCR

| | |
|---|---|
| human_ACTB_F | TCCCTGGAGAAGAGCTACG |
| human_ACTB_R | GTAGTTTCGTGGATGCCACA |
| human_UPF1_F | AGATCACGGCACAGCAGAT |
| human_UPF1_R | TGGCAGAAGGGTTTTCCTT |
| Gl3'_F | AGAAGGTGGTGGCTGGAG |
| MS2v5_R | CCGTTTGTAGGTTACCGG |
| PP7ORFv5_R | AAGAAGTGGATCCCATACC |
| Gl_Int2_F | GATGGTTCTTCCATATTCCC |
| Gl_Ex2/3_F | AGAACTTCAGGCTCCTGG |
| BAG1_F | GTGAACCAGTTGTCCAAGACCTG |
| BAG1_R | CAAGTGCTGACAACGGTGTTTCC |
| ANTXR1_F | ATGCCTTGTGGGTCCTACTG |
| ANTXR1_R | GAGGTGTGGTAGGCGTTGTT |
| GADD45A_F | GGAGGAATTCTCGGCTGGAG |
| GADD45A_R | CGTTATCGGGGTCGACGTT |
| ChIP_5'_NheI_F | CAGACACCATACTGCGGCTAGC |
| ChIP_5'_SacI_F | CACGAGCTCACCGGTG |
| Gl_Int1_R | TAACCTTGATACCAACC |

*For 3′ detection in the ChIP experiments, the primer pairs Gl3′_F and MS2v5_R or PP7ORFv5_R were used.

**Movie S1. Real-time imaging of transcription sites in WW (separate file)**

Live imaging of the U2OS cell expressing WW. The transcriptions of reporter genes were induced using Ponasterone A. The wild-type β-globin transcript expressing from either direction of promoter contains MS2 or PP7 sequences in the 3'UTR that was labeled with stdMCP-stdGFP or stdPCP-stdmScarlet as described in Figure 1. Each transcription site was imaged every 2 min.

**Movie S2. Real-time imaging of transcription sites in WW (separate file)**

Live imaging of the U2OS cell expressing WP. The transcriptions of reporter genes were induced using Ponasterone A. The wild-type or PTC-containing β-globin transcripts expressing from either direction of promoter contained MS2 or PP7 sequences in the 3’UTR that were labeled with stdMCP-stdGFP or stdPCP-stdmScarlet as described in Figure 1. Each transcription site was imaged every 2 min.
